# Emergent oscillations during cellular directional decision-making on junctions

**DOI:** 10.1101/2022.10.14.512239

**Authors:** Jonathan E. Ron, Michele Crestani, Johan M. Kux, Jiayi Liu, Nabil Al-Dam, Pascale Monzo, Nils C. Gauthier, Pablo J. Sáez, Nir S. Gov

## Abstract

Motile cells inside living tissues often encounter junctions, where their path branches into several alternative directions of migration. We present a theoretical model of cellular polarization for cells migrating along one-dimensional lines, arriving at a symmetric Y-junction and extending protrusions along the different paths that emanate from the junction. The model predicts the spontaneous emergence of deterministic oscillations between competing protrusions, whereby the cellular polarization and growth alternates between the competing protrusions. The oscillations are modified by cellular noise, but remain as a dominant feature which affects the time it takes the cell to migrate across the junction. These predicted oscillations in the cellular polarization during the directional decision making process at the junction are found experimentally for two different cell types, non-cancerous endothelial and cancerous glioma cells, migrating on patterned network of thin adhesive lanes with junctions.

## INTRODUCTION

Directed cell movement is central to processes such as organ development, tissue regeneration, cancer metastasis, and immune response [1–3]. A moving cell must negotiate obstacles that frequently generate spatial junctions or bifurcations, and upon encounter the cell has to decide on a new direction of motion [4–8]. While previous works of cells which encounter bifurcations have focused on directed migration, where the arms in the junction had a bias in hydraulic pressure [4, 6, 7], chemical cues [6], or size [5], less is known about how a single cell behaves when encountering a spatial bifurcation where the paths of the cell are symmetric, such that the cell is not directed by an external polarity cue.

While migrating inside tissues, cells navigate through complex geometries and have highly branched shapes along complex trajectories [9, 10]. Migrating cells face obstacles imposed by the surrounding extracellular matrix and cells of the surrounding tissue. Cells moving in such an environment extend protrusions that probe the alternate narrow paths [11], and therefore need to efficiently choose the optimal direction of migration. When encountering a junction along the path, the cell may either get stuck, go back or continue forward along one of the available paths [12]. An example of this process shows cancerous cancerous cells (human glioma propagating cells, hGPCs) moving on a network of thin blood vessels in a mouse brain explant (Fig.1A). Glioma cells often encounter bifurcating junctions as they invade the brain along the abluminal side of the network of blood vessels, that frequently branches with Y-shaped junctions (Fig.1A) [13].

**FIG. 1:**
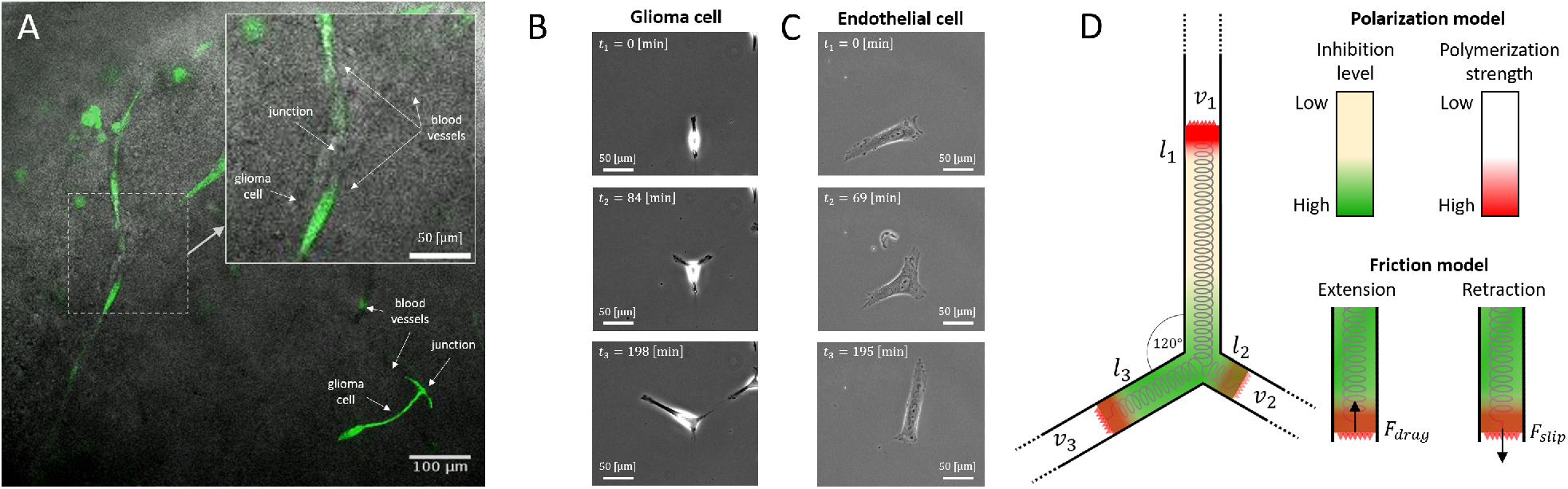
Models to study cell migration through junctions. A) Human glioma propagating cells (hGPCs) migrating in a mouse brain slice (brain slice overlay assay [24], movie 1). The cells migrate on the surface of the blood vessels in a quasi-linear motion, and encounter junctions where the blood vessels bifurcate. Upper-right inset zoom in on a bifurcating blood vessel. B) Phase contrast time-lapse images (movie 2) of an hGPC migrating along a laminin-coated Y-junction (left panels) and an endothelial cell (HUVEC) migrating along a fibronectin-coated Y-junction (right panels, movie 3). C) Model illustration. Three arms of length *l*_*i*_ with local polymerization speed *v*_*i*_ (*i* = 1, 2, 3) move along the confinement of a three-way junction (black lines). All arms are connected via a spring. Green/Yellow gradient color denote areas of low/high concentration of the polarity cue *c*(*x*) which inhibits the actin polymerization at the leading edges of the arms (red color denotes the polymerization strength). At each edge there is a friction force which acts as a simple drag force when the arm extends, and as slip-bond friction when the arm retracts [20].

In order to decipher how motile cells pass through complex paths, several *in-vitro* studies constrained cells to migrate along networks of confining one-dimensional channels or adhesive stripes [5–7, 14–18]. When moving through such a network, the cells encounter junctions, and are often observed to form multiple protrusions along the alternative paths that leave the junction. Eventually, cells have to choose one of the paths in order to leave the junction and continue their migration through the network. This migration may be random, or oriented using an external chemotactic gradient that biases the motion. Examples of such in-vitro experiments are shown in Fig.1B, where an hGPC and a HUVEC are observed to migrate across a Y-junction composed of adhesive lanes patterned over a non-adhesive background.

We present a theoretical framework that extends a model for the self-polarization of one-dimensional cells [19, 20], to include multiple competing cellular protrusions. Our simplified model of cell migration, treating the cell as composed of linear segments, allows us to obtain deep understanding of the resulting complex dynamics without describing general cellular shapes [21]. Our model predicts that during the passage of the cell over the symmetric Y-junction, the polarization of the cell undergoes *deterministic* oscillations between the competing cellular protrusions, resulting in cycles of elongation and retraction. These oscillations are an inevitable result of the polarization mechanism of the cell, which in our model is sensitive to the length changes of the competing protrusions. We find that the deterministic oscillations are modified by the inherent noise of the intracellular actin dynamics. These theoretical predictions motivated experimental studies in two different cell systems, non-cancerous endothelial cells (HUVECs) and cancerous cells (hGPCs). Both of these cell types face junctions while migrating over blood vessels during angiogenesis and repair [22], or metastasis [23]. In addition, endothelial cells and cancer cells present different migratory behavior which implies different actin polymerization regimes related to the speed of migration. We find that despite these differences, both of these two cell types display oscillatory dynamics, and exhibit motility modes along the junction which fit the different dynamical phases predicted by the model.

## THE MODEL

We extend the model of a migrating cell along a linear track [20], to the case of a cell moving over a symmetric Y-junction. When spanning the junction, the cell has three arms which are free to extend or retract (Fig.1C). Within the model, the dynamics of each arm (denoted by *i*) is described by three dynamical variables: 1) its length *l*_*i*_, 2) the local actin-polymerization speed *v*_*i*_ at the leading edge of the arm, and 3) the concentration of adhesions *n*_*i*_, at the arm’s leading edge

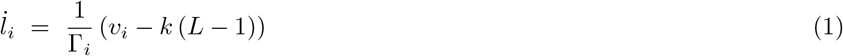

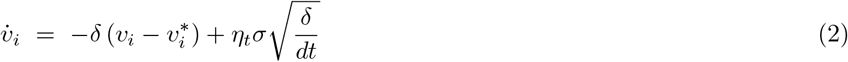

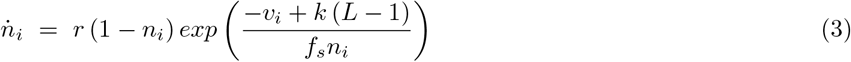

where *k* is the mean elastic constant of the cell (of rest length 1), *κ* and *f*_*s*_ are the elastic constant and force susceptibility of the adhesion molecules (which behave as slip-bonds), *δ* is the rate at which the local actin flow changes, *r* is the equilibrium dissociation constant of the adhesion molecules describing the effective adhesiveness of the surface and *L* = Σ_*j*_ *l*_*j*_ is the total length of the cell. The cellular noise is added to the equations for the actin-polymerization speeds as a random Gaussian noise (*η*_*t*_) with amplitude *σ*.

The friction term Γ_*i*_ in (Eq.1) describes a constant drag when an arm extends, and slip-bond adhesion friction when an arm retracts

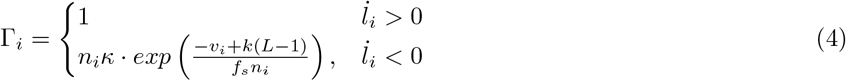

This allows us to describe stick-slip dynamics of the cell [20].

In the model, the net global actin treadmilling flow within an arm, *u*_*i*_, advects a polarity cue which inhibits the local actin-polymerization (or local flow) *v*_*i*_ emanating from the leading edges of the arms (Fig.1C). The steady-state global and local flows, as function of the arms lengths and the local concantration of the polarity cue, are given by

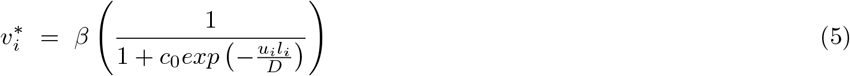

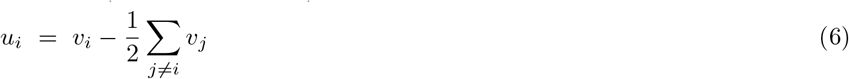

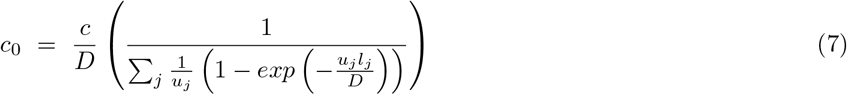

where *D* is the diffusion coefficient of the polarity cue, and *c* is a dimensionless quantity which represents the ratio between the total and saturation concentrations of the polarity cue. A key parameter in our model is *β*, denoting the maximal actin polymerization speed produced at the leading edge of the cellular arms.

The steady-state local actin flow (Eq.5) describes the inhibitory effect of the polarity cue on the local actin-polymerization activity, at each arm. The actin flow from each arm is assumed to split evenly at the junction, and therefore contributes a half of its flux when calculating the net flow within in each arm (Eq.6). It is this net flow that advects the polarity cue, which explains why the exponential functions that describe the polarity cue at the edges of the arms, contains these net flow speeds (Eqs.5,7). The total amount of polarity cue, its integral over the entire cell, is conserved. For the full solution of the polarity cue distribution along a cell with *N* protruding arms see Supplementary Information S1.

## SYMMETRY BREAKING, POLARIZATION AND EMERGENT OSCILLATIONS

Within our model, cell migration occurs when the cell spontaneously self-polarizes, due to the feedback between the actin flow and the intracellular distribution of a polarity cue: The polarity cue is advected by this flow, and in turn determines the flow by inhibiting actin-polymerization at the leading edges of the arms (Eq.5). We analyze the conditions that allow cells to polarize and become migratory on the Y-junction, for two limiting cases: 1) A symmetric case, in which the cell lands at the center of the junction and spreads symmetrically in all direction (Fig.2A), and 2) a moving case, in which a cell which migrates at steady-state along a linear track [20] enters the junction, and grows two symmetric protrusions along the new arms (Fig.2D). The first case is simpler to solve analytically but is not often observed experimentally, while the second case is the more common experimental observation.

**FIG. 2:**
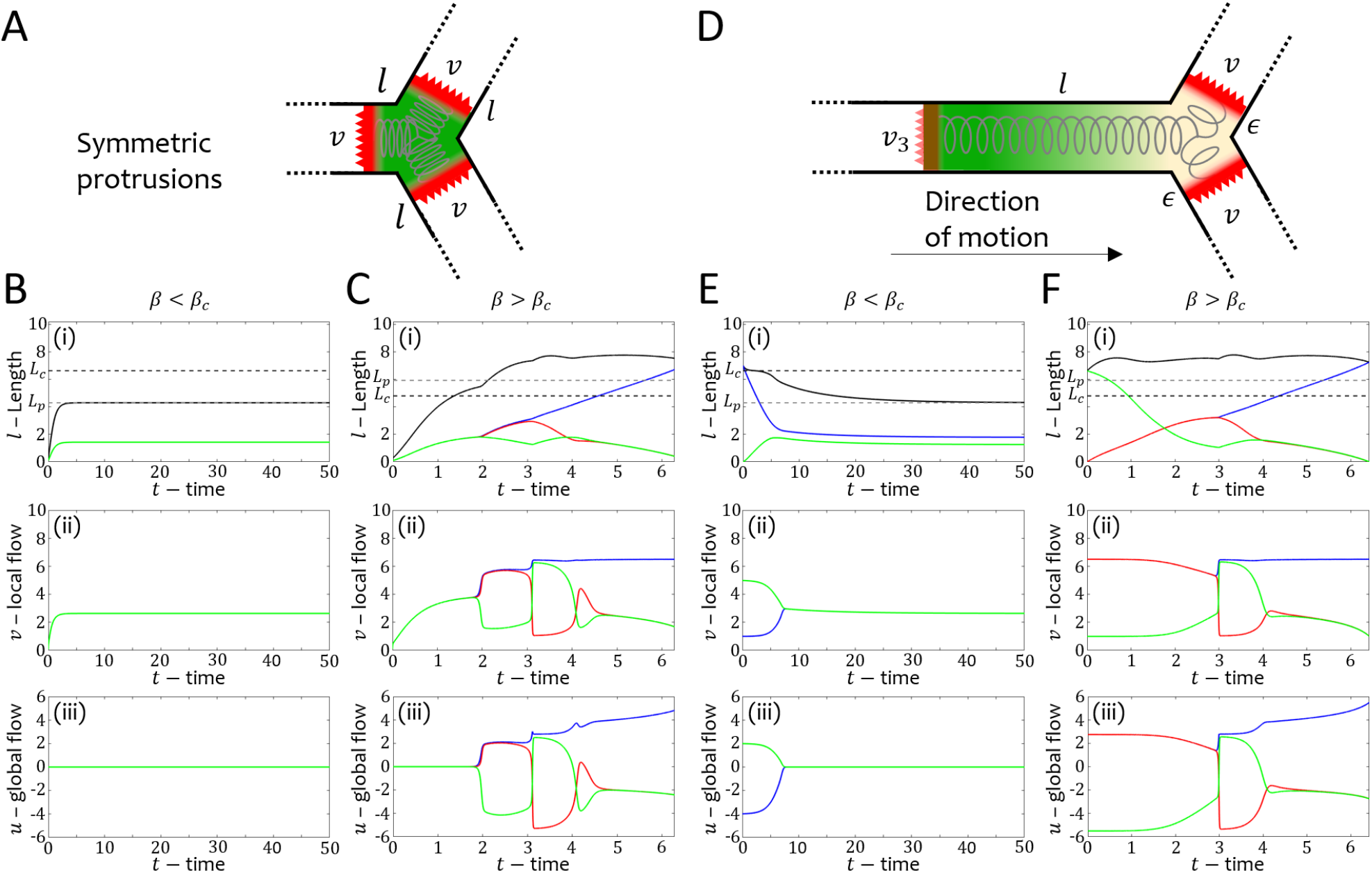
Deterministic oscillations control directional decision making. A) The symmetric spreading case, where all the arms initially extend symmetrically, with the same local actin-polymerization speed *v*. B) For *β* = 5 *< β*_*c*_ the cell reaches a stationary shape, with all the arms of equal lengths and actin-polymerization activity. C) For *β* = 6.5 *> β*_*c*_ the cell polarizes and leaves the junction. i-iii) Time series of the arm lengths, actin-polymerization speeds and net global actin flows. D) The moving case: a cell of length *l* enters the junction. At the front, the cell extends two symmetric protrusions, initially of equal length (*ϵ*) and the same local actin-polymerization speed *v*. At the rear, the actin-polymerization is *v*_1_. E) For *β* = 5 *< β*_*c*_ the migrating cell stops at the junction. F) For *β* = 6.5 *> β*_*c*_ the cell migrates past the junction, leaving it through one of the new protruding arms. i-iii) Time series of the arm lengths, actin-polymerization speeds and net global actin flows. Legend: Blue/Red/Green curves represent arm 1/2/3 respectively. Dashed Black/Dashed Gray lines indicate *L*_*c*_/*L*_*p*_ respectively. Parameters: *δ* = 250, *c* = 3.85, *D* = 3.85, *k* = 0.8, *r* = 5, *f*_*s*_ = 5, *κ* = 20, *σ* = 10^−7^ (noise amplitude).

The polarization of the cell is critically dependent in our model on its length [20]. For the symmetrically spreading cell (Fig.2A), we can calculate the length (*L*_*p*_) where there is a balance between the protrusive forces and the elasticity of the cell [20], given by

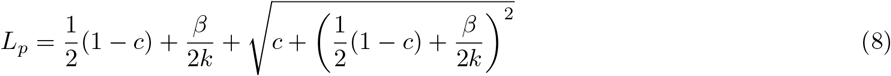

The critical length *L*_*c*_ describes the length in which the feedback between the treadmilling flow and the polarity cue destabilizes the uniform solution, and the cell becomes polarized

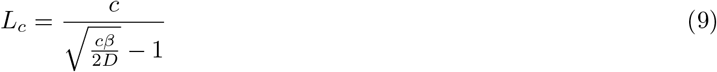

By equating (Eqs.8,9) we obtain the critical polarization speed

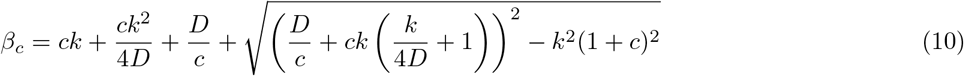

For *β < β*_*c*_ the cell spreads and reaches a symmetric, stationary shape that is stable on the junction (Fig.2B). For *β > β*_*c*_ the cell spreads and spontaneously polarizes along one of the arms, when its total length exceeds *L*_*c*_ (Fig.2C). The cell eventually migrates away from the junction, as indicated by two of the arms shrinking to zero (Fig.2C). For the moving cell (Fig.2D) we find that symmetry breaking occurs for slightly smaller values of *β* compared to the symmetric spreading case (Fig.2E,F). Below a critical value of *β* the moving cell stops at the junction, and remains stationary (Fig.2E), while above this critical value the cell is able to continue migrating across the junction (Fig.2F). We provide further approximations for the critical length in the case of the moving cell in Supplementary Information section S2.

Note that *β*_*c*_ in (Eq.10) is larger than the critical value for spontaneous motion along a straight line (*β*_0_). This is due to the additional leading edge that the cell has on the Y-junction, which increases the dilution of the polarity cue, which is the limited resource that is needed for spontaneous polarization. Therefore we predict that a cell with *β*_*c*_ *> β > β*_0_ will get trapped in the junction, although *β* is sufficiently large for one-dimensional migration.

In both cases, of symmetric spreading and migration to the junction, above the critical *β* there are clear oscillations in the local actin-polymerization activity at the tips of the competing arms (Fig.2C,F), as the cell moves past the junction. These oscillations in the actin-polymerization activity affects the growth and retraction of the arms, as seen in their length dynamics. These are *deterministic oscillations*, not driven by noise that is negligible in these simulations.

The origin of these oscillations is due to the redistribution of the conserved amount of polarity cue over the entire length of the cell, which is highly sensitive to the length changes. We analyze the origin of these oscillations in more detail for a symmetrically spreading cell (Fig.3), which undergoes the symmetry-breaking event (*L > L*_*c*_) at *t ∼*2. At this time one arm “loses”, as indicated by its decreasing polymerization speed (green line in Fig.3A(iii)), while the two remaining arms compete for the cell polarization direction (blue and red lines in Fig.3A(iii)). There are then two oscillation cycles, denoted by the dashed vertical lines in Fig.3A. During these oscillations the actin-polymerization speeds (*v*_*i*_) undergo rapid and large changes (Fig.3A(iii)), driving corresponding changes to the lengths of the arms (Fig.3A(i)).

**FIG. 3:**
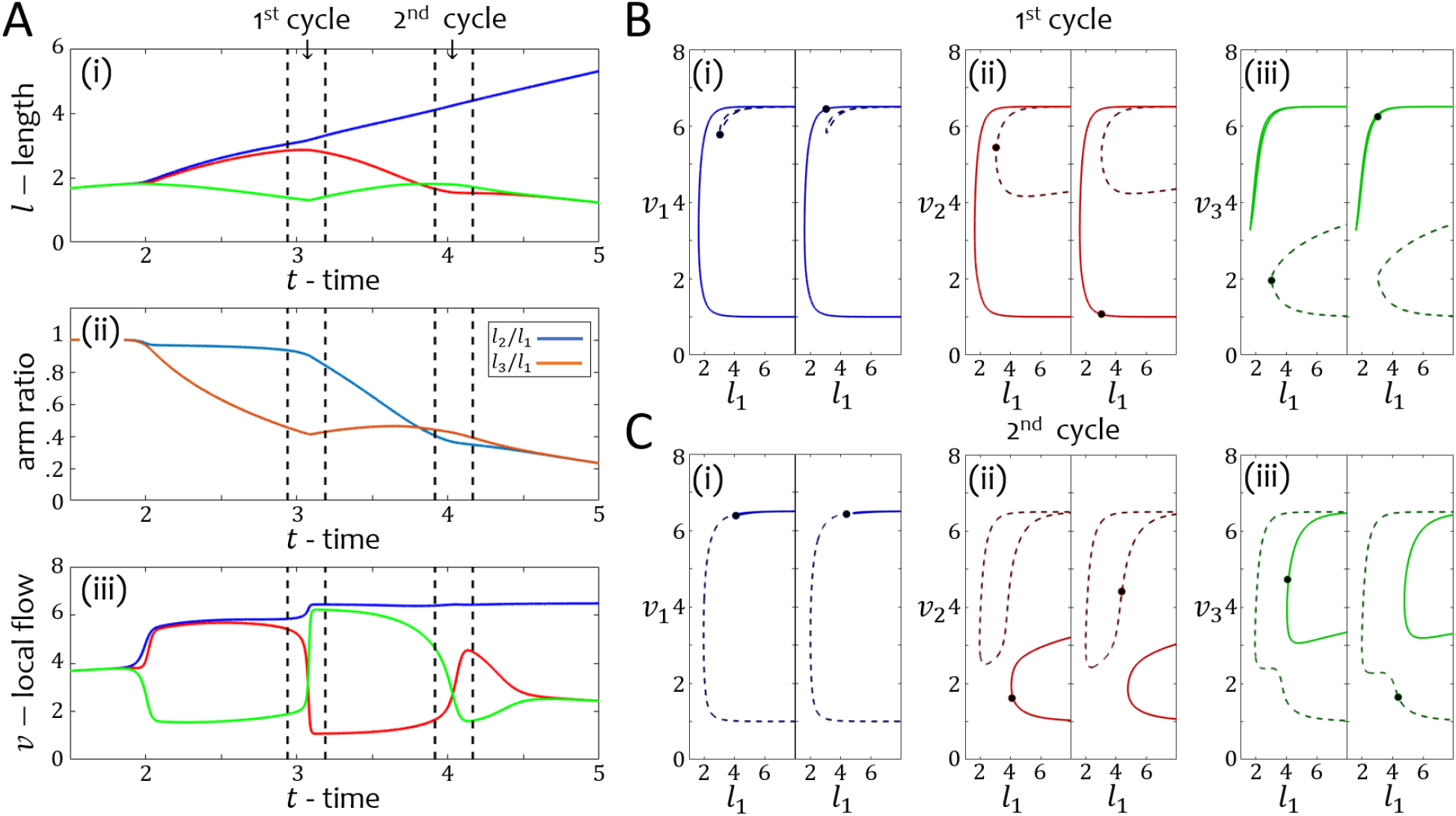
Oscillation analysis of the symmetric case. A) Time series of the (i) length, (ii) the arm ratio and (iii) the local flow respectively. Red/Blue/Green colors in (i,iii) correspond to arms 1/2/3 respectively. Blue and Orange curves in (ii) correspond to the arm ratios *l*_2_*/l*_1_ and *l*_3_*/l*_1_ respectively. Black dashed lines indicate the examined cross-sections of the first and second cycles of oscillations. B-C) The solutions branches of the local flows in the cross-sections of the first and second cycle respectively. (i)/(ii)/(iii) correspond to *v*_1_/*v*_2_/*v*_3_. Left/Right panels correspond to the solutions branches before/after the transition. Points indicate the values from the simulations in (A,iii). Parameters: *δ* = 250, *β* = 6.5, *c* = 3.85, *D* = 3.85, *k* = 0.8, *f*_*s*_ = 5, *r* = 5, *κ* = 20, *σ* = 10^−7^ (noise amplitude).

We analyze the oscillatory mechanism by plotting the steady-state solutions of the actin-polymerization speeds in the proximity of the oscillations (Fig.3B-C). For each oscillation, we chose a time just before and after the oscillation cycle (vertical dashed lines in Fig.3A), and use the simulations to calculate the ratio between the lengths of the arms at these times (Fig.3). We then plot the self-consistent solutions for the actin-polymerization speeds (Eq.5) as a function of the length of the winning arm (*l*_1_), while maintaining the ratios fixed (Fig.3B-C). The solutions associated with the value of *l*_1_ from the simulations are denoted by the circle marker (Fig.3B-C). The steady-state solutions are obtained using a continuation method [25].

This analysis shows that the mechanism for the deterministic oscillations lies in transitions between the different solution branches as the arms elongate/retract and the ratios of their lengths change. Just before the oscillations we find that the solutions of the losing arms (*v*_2_/*v*_3_, red/green respectively) are at the tips of a solution branch saddle (Fig.3B(ii,iii),C(ii,iii)). As *l*_1_ grows larger, the ratios between *l*_1_ and the losing arms *l*_2_ and *l*_3_ decreases (Fig.3A(ii)), and as a result the saddle shifts slightly, yet sufficiently to force the solutions to discontinuously switch between the different solution branches. These branch transitions are most significant for *v*_2_ and *v*_3_, but also occurs for *v*_1_ in the first oscillation (Fig.3B(i)).

We emphasize that these non-linear deterministic oscillations, do not arise due to growing arm(s) simply pulling on the remaining arms, causing them to retract. Rather, the oscillations arise due the sensitive inter-dependence of the distribution of the finite amount of polarity cue over the whole cell, and its effect on the local actin polymerization activity at the ends of the competing arms. Due to this competition, the actin-polymerization activity shifts between the leading edges of the competing arms, such that they oscillates out-of-phase with each other. Using the same analysis, we also find another oscillation pattern, of smaller amplitude, which is associated with the stabilization of the flows in the losing arms (Supplementary Information section S3).

In Fig.4 we show examples of HUVEC and hGPC cells performing the out-of-phase oscillations in the local actin polymerization activity at the tips of the arms, predicted by the model (for the experimental methods see Supplementary Information section S4). We quantify the actin polymerization activity at the tips of the arms in the experiments as follows: we define a region extending over a distance of 10 microns back from the edge of each arm. Within this region, we measure the overall membrane area (“tip area”), which captures the amount of protrusive lamellipodia activity. We also measure the total actin fluorescence intensity (“tip actin intensity”) within this region, as a measure for the actin polymerization activity at the leading edge of the arm. Both of these measures are compared to the local actin polymerization speed in the model (*v*_*i*_, Eq.2), and both exhibited the predicted out-of-phase oscillations in the losing arms (Fig.4A(ii,iii),E(ii,iii)).

**FIG. 4:**
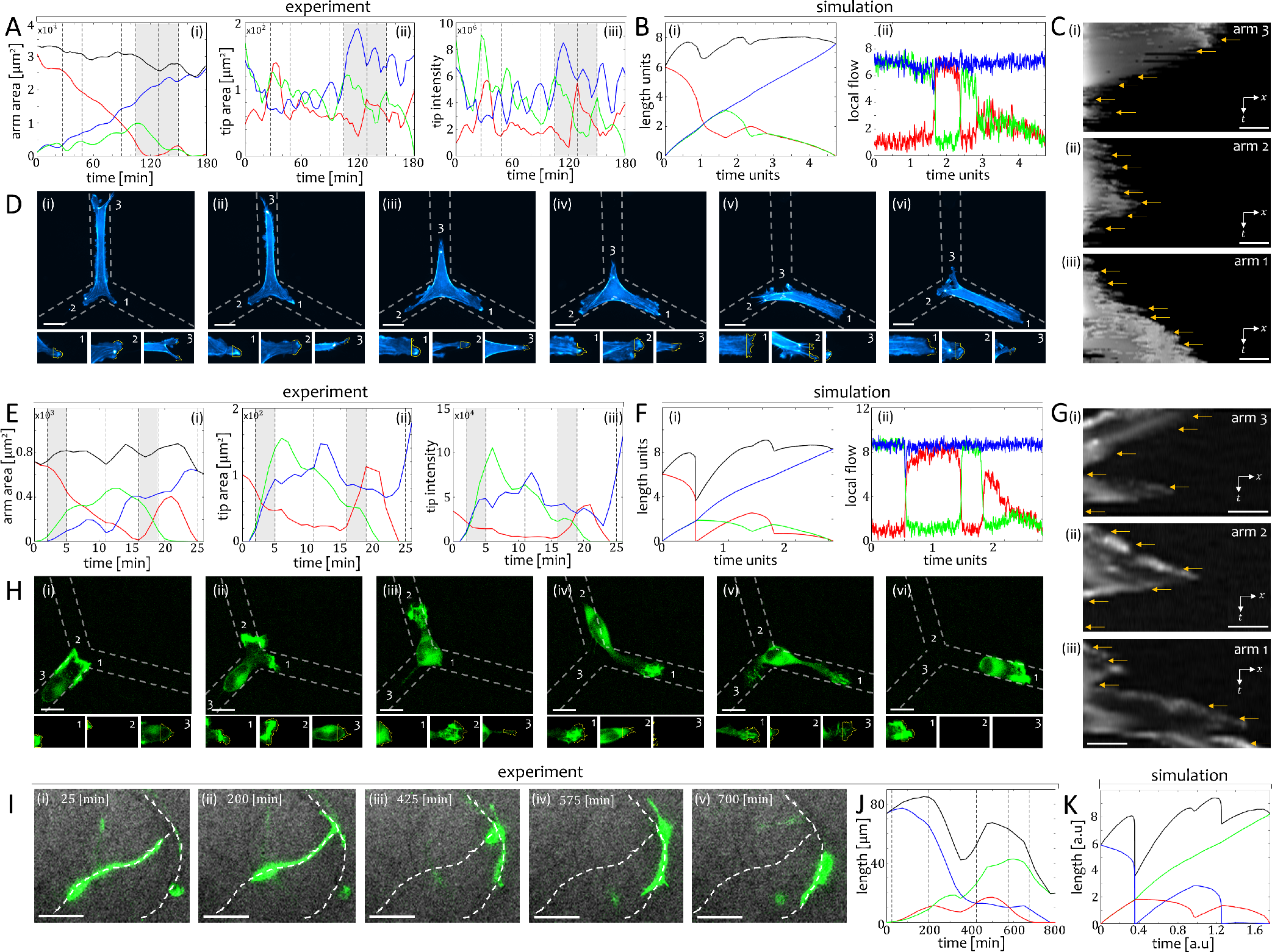
A) HUVEC experiment. i) Total area of the arms. ii) Lamellipodia (arm tip) area. iii) Lamellipodia actin intensity. B) Simulation with (*β, σ*) = (6.7, 0.5). i) Arm length. ii) local actin flows. Blue/Green/Red indicate arm 1/2/3. Black indicate the total cell length. Gray section indicate the regions where the actin flow flips between the losing arms. Gray dashed lines correspond to the images in D and the arrow markings in C. C) Arm kymographs. D) Time-lapse images (movie 4). Upper panels: Whole cell. Lower panels: Enlarged arm protrusions. Yellow is a 10 [µm] window from which the intensity was measured. E) hGPC experiment. i) Total arm area. ii) Lamellipodia (arm tip) area. iii) Lamellipodia actin intensity. F) Simulation with (*β, σ*) = (8.7, 0.3). i) Arm length. ii) local actin flows. Blue/Green/Red indicate arm 1/2/3. Black indicate the total cell length. Gray section indicate the regions where the actin flow flips between the losing arms. Gray dashed lines correspond to the images in H and the arrow markings in G. G) Arm kymographs. H) Time-lapse images (movie 5). Upper panels: Whole cell. Lower panels: Enlarged arm protrusions. Yellow is a 10 [µm] window from which the intensity was measured. I) Time-lapse images of an hPGC cell migrating in a mouse brain slice (movie 6). White dashed lines indicate the blood vessels over which the cell migrates upon. J) Experimental time-series of the lengths of the cellular arms. Dashed vertical lines correspond to the images in I. K) Time-series of the arms’ lengths in simulation, using (*β, σ*) = (10.5, 0.5). Scale bars in (A-G) are 100 [µm]. Scale bars in (I) are 50 [µm]. Simulation parameters: *c* = 3.85, *D* = 3.85, *k* = 0.8, *f*_*s*_ = 5, *r* = 5, *κ* = 20, *δ* = 250.

We also present an example of an hGPC migrating along blood vessels, which contains bifurcations in an *ex-vivo* mouse brain slice (Fig.4I,J), and compare it to simulations (Fig.4K). We find the same oscillatory dynamics observed on the patterned lanes in Fig.4A-G, suggesting the validity of our model for cellular dynamics in the *in-vivo* physiology. For more comparisons of the oscillations observed in experiments with the model see Supplementary Information section S5 (Figs.S3,S4). In Fig.S4A-D we provide an example of a cell which resides on the center of the junction and exhibits symmetric spreading along all the available arms as in the theoretical symmetric case (FIG 2A).

## ESCAPE TIMES AND TRAPPING AT THE JUNCTION

Next, we investigate the dependence of the average time it takes the cell to migrate past the junction (“escape time”, *T*_*escape*_) on the actin-polymerization activity (*β*), for a finite value of the actin response rate *δ* (Eq.2) (the case of *δ* → *∞*, in which the actin flow instantaneously returns to its steady-state, is shown in the Supplementary Information section S6). For the moving cell, the escape time is defined as the time from when the cell first arrives, to when it leaves the junction. For both the symmetric spreading and the moving cell (red and black lines respectively in Fig.5A-C), we observe the same trends: First, below the critical value of *β*_*c*_ the cell gets trapped on the junction, and the escape time diverges. As *β* increases above this threshold value, the cell migrates past the junction faster (Fig.5A). This is expected, since the cell migration speed increases with increasing *β*. However, above a critical value of *β*, there appears a new “slow” dynamical process, with increasing probability (Fig.5C). The transition to this slow process occurs at a higher value of *β* in the case of the moving cell (Fig.5C).

**FIG. 5:**
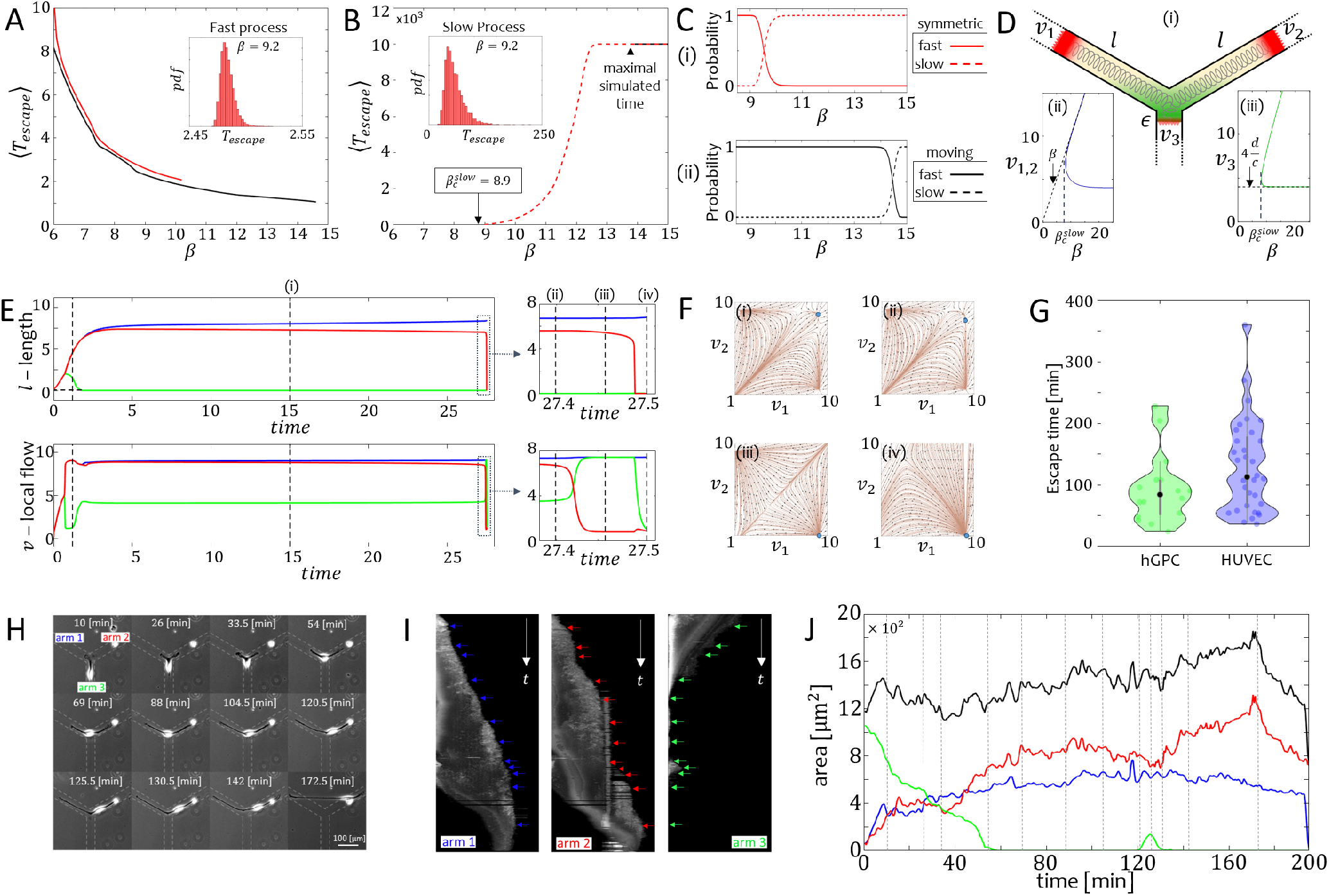
Mean escape time from the junction. A) The mean escape time as a function of *β*, for a symmetric cell spreading on the junction (red) and a migrating cell (black). The dynamics in this regime corresponds to the processes shown in Figs.2,4. In (B) we plot the mean escape times for the “slow” process (dashed lines), which is described in (E-G). Insets of (A,B) give the escape time distributions for both the fast and slow processes at *β* = 9.2. C) The probability to be in the meta-stable slow process (dashed lines) or in the fast process (solid lines), for the symmetric case (red) and the moving case (black) as a function of *β*. D) The limit of the theoretical model where two arms compete and balance the spring force. (i) Illustration of the model where two arms balance the spring force: *v*_1,2_ is the local flow in the long arms 1, 2, where *v*_1_ = *v*_2_. *l* is the arm length of arms 1 and 2 respectively (*l*_1_ = *l*_2_), and *v*_3_*/ϵ* is the local flow/length of arm 3. Green gradient denotes the concentration of the inhibitor of actin-polymerization, while red color denotes the actin-polymerization strength at the tips of the arms. (ii)/(iii) The steady-state solutions for *v*_1*/*2_/*v*_3_ which correspond to the illustration in panel (i) (Eq.13). Black dashed lines indicate the steady-state solutions in the limit of *l* → *∞* (Eqs.14,15), and the saddle node bifurcation at the threshold value of *β* is denoted. E) Time series of the length (upper panels) and local actin flow (lower panels) for the slow process (*β* = 9.2) in the symmetric case. Right insets zoom in on the final stage, where the balance between the long arms is resolved. Blue/Red/Green indicate arm 1/2/3. F) The theoretical steady-state *v*_1_-*v*_2_ flow fields for the cross-sections (i-iv) in (E). Blue point indicates the values of *v*_1*/*2*/*3_ values taken from the simulation at each of these times, showing that the slow process corresponds to a local meta-stable solution, which eventually becomes unstable. G) Violin bar plot of the escape times of hGPC cells (green) and HUVEC cells (blue). H) Time-lapse images (movie 7) which correspond to times indicated by arrows in the kymographs (I), and by the black dashed lines in the length time series (J). Blue/Red/Green indicate arm 1/2/3. Black indicates the total length of the cell. Simulation Parameters: *c* = 3.85, *D* = 3.85, *k* = 0.8, *f*_*s*_ = 5, *r* = 5, *κ* = 20, *δ* = 250.

The slow process is characterized by two arms which extend symmetrically with very similar lengths (Fig.5D,E), while the third arm shrinks to zero length. Eventually the competition between the long arms is resolved by an oscillation in the actin polymerization activity in one of the long arms (Fig.5E), resulting in its collapse (Fig.5E,F).

We can estimate the threshold value, 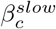, above which the slow process emerges by taking the limit where two arms of equal lengths *l* balance the spring force (Fig.5D), i.e, *v*_2_ = *v*_1_ *≡ v*

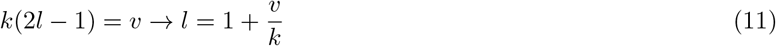

For this limit, the steady-state local flows are given by

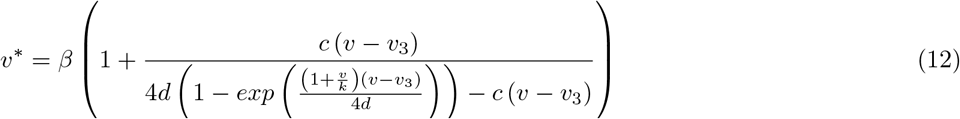

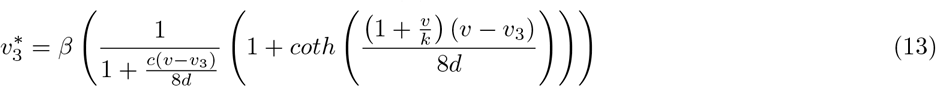

where in the limit of large *l* for *v*_1*/*2_ and small *l* for *v*_3_ (Eq.5) we obtain

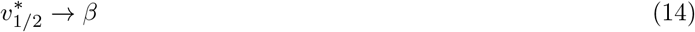

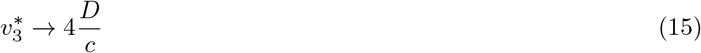

These solutions are indicated in Fig.5D(ii,iii), and the saddle node bifurcation at the threshold value of *β* which is found to be in the proximity of the value that we find in the simulations for the spreading cell (Fig.5D,E). In Fig.5F we show that the slow mode is formed due to a local stable solution with *v*_1_ *≃ v*_2_, which eventually becomes unstable. Further analysis of the slow process is given in the Supplementary Information section S7.

When comparing these cell types (Fig.4), we found that the cancerous hGPC correspond to a higher value of the actin polymerization parameter *β*, compared to the non-cancerous HUVEC. In agreement with this conclusion, the hGPCs have shorter escape times past the junction, compared to HUVECs (Fig.5G). In addition, hGPCs exhibit more often events that correspond to the slow process predicted by the model for cells with higher values of *β* (Fig.5H-J). Note that in the experiments some of these events are resolved when the cell detaches from the junction and remains stretched over the non-adhesive region between the two competing arms. This detachment is beyond the model.

The escape time also depends on the amplitude of the noise in the local actin polymerization activity *σ* (Eq.2). This is demonstrated in Fig.6A for a moving cell in the stick-slip migration regime (*β* = 8). For a low noise level (Fig.6A(i)) the escape time is sharply defined, and the cell always leaves the junction along either one of the new arms.

**FIG. 6:**
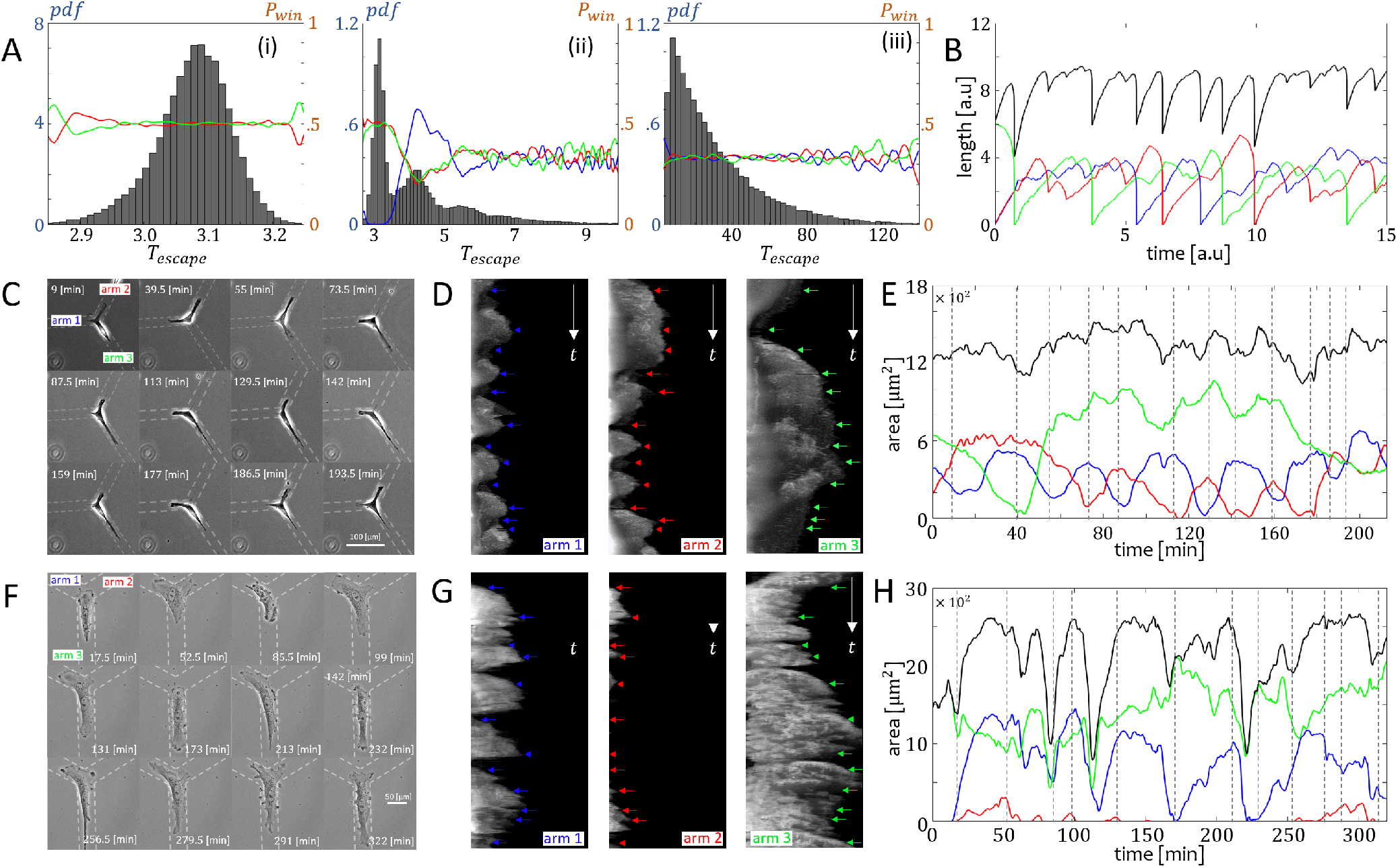
Noise effects on the dynamics at the junction. A) The escape time probability density function (left y-axis in blue) and the probability for the cell to leave the junction along each of the arms (right y-axis in orange), for noise amplitudes (Eq.2): *σ* = 0.4 (i),*σ* = 1.2 (ii), *σ* = 1.6 (iii). Blue/Red/Green indicate arm 1/2/3, respectively. B) Example of a simulation length time series with *σ* = 1.6. C-D) hGPC and F-G) HUVEC experiments showing cells that are trapped on the junction. Left panels (C,F): Time-lapse images (movies 8 and 9). Middle panels: Arm kymographs. Blue/Red/Green arrows indicate the time stamps which correspond to the time-lapse images. Right panels (E,H): Experimental time series of the area of the cellular arms. Gray dashed lines correspond to the time-lapse images. Blue/Red/Green indicate arm 1/2/3. Black indicate the total cell length. Parameters: *β,c* = 3.85, *D* = 3.85, *k* = 0.8, *f*_*s*_ = 5, *r* = 5, *κ* = 20, *δ* = 250.

As the noise amplitude increases, the escape time increases (Fig.6A(ii)). In particular, we find that the noise gives rise to additional oscillations during the migration past the junction. Each additional oscillation increases the escape time, which has several peaks. We find that a single additional oscillation reverses the direction in which the cell leaves the junction, increasing its probability to “reflect” (leave along the direction from which it has arrived). For a larger number of oscillations the cell has equal probability to leave along any of the three arms. This is also observed for a high noise level (Fig.6A(iii)), where any memory of the original direction of motion is lost, and the escape time distribution becomes exponential. In this limit, we often observe cells that are “stuck” on the junction, for very long times (Fig.6B).

The increased trapping of the cell on the junction at higher noise is unusual at first sight, as noise is normally attributed with triggering the escape of particles from traps [26, 27]. However, for the cell to leave the junction it needs to maintain its polarity for the duration of the escape time, and the large noise diminishes the persistence time of the polarization, inhibiting the escape probability. Nevertheless, the noise also plays a role in aiding to the cell’s escape, whereby during the slow process the meta-stable configuration is destabilized (Fig.5).

In Fig.6 we provide an example of an hGPC and a HUVEC which exhibit many cycles of elongations and retractions of the arms, unable to maintain a persistent polarization, and therefore remain trapped on the junction. Both examples correspond to the large noise limit of our model (Fig.6B). Further analysis of the effects of noise on the dynamics at the junction is given in the Supplementary Information section S8 (Figs.S5,S6). In Fig.S5E-H, we show an example of a cell that “reflects” at the junction (leaves along the direction from which it arrived), as occurs in our model when the actin flow is subjected to intermediate values of noise. (Fig.6A(ii)).

## DISCUSSION

We have presented here a new theoretical model for the directional decision making process at the level of a single migrating cell, when moving past a symmetric Y-junction. Our model describes the spontaneous symmetry breaking during cellular spreading over a junction, and predicts *deterministic* oscillations during the migration of a cell past a junction. These oscillations arise due to the competition for the finite resource, the polarity cue, which is redistributed over the arms as their lengths evolve. As a result, the actin polymerization activity oscillates between the competing leading edges. These out-of-phase oscillations are most prominent between the two “losing” arms, and also induce some oscillation in the length dynamics of the cellular protrusions. This is a new form of deterministic oscillations during cell migration, distinct from oscillations induced by stick-slip adhesion [20, 28] or periodic macropinocytosis [29].

These surprising theoretical predictions are validated by experimental data acquired from two cell types, cancerous hGPCs and non-cancerous HUVECs. We find that the predicted out-of-phase oscillations in the lamellipodia activity at the leading edges of the cellular protrusions, and the protrusions’ lengths, are observed in these experiments when these cells migrate past the junction. These comparisons expose that the non-cancerous (HUVECs) and cancerous cells (hGPCs), correspond to low/high levels of motility regimes, respectively.

The model predicts that cells with low actin polymerization activity will be prone to remain trapped on junctions, even if they are well polarized and motile when moving along simple linear tracks. As expected, the mean trapping time on the junction decreases as the cells become more motile (higher actin polymerization activity). Larger noise in the actin polymerization activity acts to increase the number of polarization oscillations and the overall trapping time at the junction. Interestingly, we find that when the polymerization activity is high, the cells may be transiently trapped on the junction, due to the competition between two symmetric and highly elongated protrusions. The latter suggest that efficient migration across a complex geometry and resolution of directional dilemma occurs within a range of actin polymerization that is sufficiently strong to allow motility, but that is weak enough to prevent the non-productive consequence of excessive acto-myosin interaction. These predictions will be the subject of future experimental studies. Future extensions of the model might include asymmetric junctions [8], different geometries and the interaction with other cells to evaluate the effect of cell-to-cell interaction[30, 31]. In addition, this model could be used to explore the effects of chemotaxis and barotaxis on the cellular decision-making and migration.

To conclude, we have presented a detailed, yet simple, model for the mechanism of cellular directional decision making on a Y-junction. In this model the decision is performed due to spontaneous symmetry breaking, involving deterministic oscillations. The model explains how the duration of the directional decision making depends on the internal motility parameters of the cell, offering a natural explanation for the puzzling slowness of this process in many cells [18].

## ACKNOWLEDGMENTS

J.E.R. and N.S.G. acknowledge useful discussions with Raphael Voituriéz and Arik Yochelis. N.S.G. is the incumbent of the Lee and William Abramowitz Professorial Chair of Biophysics, and acknowledges support by the Israel Science Foundation (Grant No. 1459/17). This research is made possible in part by the historic generosity of the Harold Perlman Family. This work was supported by the Human Frontier Science Program (HFSP) RGP0032-2022 to P.J.S. and N.S.G., by the Deutsche Forschungsgemeinschaft (DFG) grant number 335447717-SFB 1328 (project A20), and Forschungszentrum Medizintechnik Hamburg (FMTHH) 04fmthh2021 to P.J.S. This work was supported by IFOM (starting package), the Mechanobiology Institute of Singapore (grant WBSR-714-016-007 -271), and the Italian Association for Cancer Research (AIRC) (Investigator Grant [IG] 20716 to N.C.G. and doctoral fellowship 3-year fellowship “MilanoMarathon-oggicorroperAIRC” - Rif. 22461 to M.C.). M.C. is a PhD student within the European School of Molecular Medicine (SEMM).

