## Supplementary Information file for "Emergent oscillations during cellular directional decision-making on junctions"

Jonathan E. Ron[1], Michele Crestani[2], Johan M. Kux[3], Jiayi Liu[1,4], Nabil Al-Dam[3], Pascale Monzo[2], Nils C. Gauthier[2], Pablo J. Sáez[3] and Nir S. Gov[1]
[1]*Department of Chemical and Biological Physics, Weizmann Institute of Science, Israel*
[2]*IFOM, FIRC Institute of Molecular Oncology, Milan, 20139, Italy*
[3]*Cell Communication and Migration Laboratory, Institute of Biochemistry and Molecular Cell Biology, Center for Experimental Medicine, University Medical Center Hamburg-Eppendorf, Hamburg, Germany and*
[4]*Zhejiang University, Hangzhou, China*

### S-1. UCSP RELATIONS FOR A CELL ON AN N-ARM JUNCTION

Consider a system of a cell on a junction with $N$ arms which are equally spaced.

The local actin flow (or polymerization force) in arm $i$

$$v_i = \beta \left( \frac{1}{1 + \frac{c}{D}\left( \frac{exp\left(-\frac{u_i x_i}{D}\right)}{\sum_j \frac{1}{u_j}\left(1 - exp\left(-\frac{u_j x_j}{D}\right)\right)} \right)} \right) \tag{S-1}$$

where $c$ is a dimensionless unit which determines the ratio between the total concentration and saturation of the polarity cue in the cell ($c = \frac{c_{tot}}{c_s}$), $D$ is the diffusion coefficient of the polarity cue, $\beta$ is the upper bound, or maximal treadmilling flow in, and $u_i$ is the global net flow with respect to each arm $i$ which is given by

$$u_i = v_i - \sum_{j \neq i} \frac{v_j}{N-1} \tag{S-2}$$

The global flow at each segment is given by the advection-diffusion of a polarity cue $c(x)$ along the segment

$$\frac{\partial}{\partial x}\left( u_i c_i(x) + D\frac{\partial c_i(x)}{\partial x} \right) \longrightarrow c_i(x) = A_0 exp\left(-\frac{u_i x}{D}\right) + B_0 \tag{S-3}$$

By applying a no-flux boundary condition at the edge of the cell (on each arm) we obtain

$$u_i c_i(x) + D\left.\frac{\partial c_i(x)}{\partial x}\right|_{x=l} = 0 \longrightarrow B_0 = 0 \tag{S-4}$$

and by considering mass conservation of the polarity cue

$$c_{tot} = A_0 \sum_i \int_0^{x_i} exp\left(-\frac{u_i x}{D} dx\right) \longrightarrow A_0 = \frac{c_{tot}}{D}\left( \sum_i \frac{1}{u_i}\left(1 - exp\left(-\frac{u_i x_i}{D}\right)\right) \right) \tag{S-5}$$

therefore we obtain that the concentration profile of the polarity cue along each arm is given by

$$c_i(x) = \frac{c_{tot}}{D} \frac{exp\left(-\frac{u_i x}{D}\right)}{\sum_j \frac{1}{u_j}\left(1 - exp\left(-\frac{u_j x_j}{D}\right)\right)} \tag{S-6}$$

Together with the continuity condition at the junction, and the conservation of the total amount of polarity cue (Eq.S-5), we solve for the polarity cue distribution along all the arms, at each instant of time.

We consider a saturation profile in the form of a Hill function for the polarity cue

$$\tilde{c}_i(x) = \frac{c_s}{c_s + c_i(x)} \tag{S-7}$$

such that the local flow (or polymerization force) is given by

$$v_i = \beta \left( \frac{1}{1 + \frac{\frac{c}{D} exp\left(-\frac{u_i x_i}{D}\right)}{\sum_j \frac{1}{u_j}\left(1 - exp\left(-\frac{u_j x_j}{D}\right)\right)}} \right) \tag{S-8}$$

where $c = \frac{c_{tot}}{c_s}$.

### S-2. THE CRITICAL LENGTHS

By combining (Eqs.S-1,S-2), the global flow can be written as

$$u_i = \beta\left(\frac{1}{1+\frac{c}{D}\left(\frac{exp\left(-\frac{u_i l_i}{D}\right)}{\sum_j \frac{1}{u_k}\left(1-exp\left(-\frac{u_k l_k}{D}\right)\right)}\right)} - \frac{1}{N-1}\sum_{j\neq i}\frac{1}{1+\frac{c}{D}\left(\frac{exp\left(-\frac{u_j l_j}{D}\right)}{\sum_j \frac{1}{u_k}\left(1-exp\left(-\frac{u_k l_k}{D}\right)\right)}\right)}\right) \tag{S-9}$$

Next, we choose small flows $u_i = \epsilon$ and $u_j = -\frac{\epsilon}{N-1}$ (as on a line, where one segment competes with the others), and also choose $l_i = l$ (equal segments)

$$\epsilon \ = \ \beta\left(\frac{1}{1+\frac{\frac{c}{D}exp\left(-\frac{\epsilon l}{D}\right)}{\frac{1}{\epsilon}\left(1-exp\left(-\frac{\epsilon l}{D}\right)\right)-\frac{(N-1)^2}{\epsilon}\left(1-exp\left(\frac{\epsilon l}{(N-1)D}\right)\right)}} - \frac{1}{1+\frac{\frac{c}{D}exp\left(\frac{\epsilon l}{(N-1)D}\right)}{\frac{1}{\epsilon}\left(1-exp\left(-\frac{\epsilon l}{D}\right)\right)-\frac{(N-1)^2}{\epsilon}\left(1-exp\left(\frac{\epsilon l}{(N-1)D}\right)\right)}}\right) \tag{S-10}$$

expand (Eq.S-10) into first order in $\epsilon$

$$\epsilon = \epsilon\left(\frac{cl^2N^2\beta}{D(n-1)(c+lN)^2}\right) + O(\epsilon)^2 \tag{S-11}$$

solve (Eq.S-11) for $l$

$$l_c(N) = \frac{1}{N}\left(\frac{c}{\sqrt{\frac{c\beta}{D(N-1)}}-1}\right) \tag{S-12}$$

and substitute $l = \frac{L}{N}$ to obtain

$$L_c(N) = \frac{c}{\sqrt{\frac{c\beta}{D(N-1)}}-1} \tag{S-13}$$

By choosing $N = 2$ we obtain the critical for a cell migrating along a linear track [1]

$$L_c(N=2) = \frac{c}{\sqrt{\frac{c\beta}{D}}-1} \tag{S-14}$$

Below the critical length, in the limit of $u \to 0$, the local flows of the arms are given by

$$v_i = \lim_{u\to 0}\left(\frac{\beta}{1+\frac{c}{D}\left(\frac{exp\left(-\frac{u_i x_i}{D}\right)}{\sum_j \frac{1}{u_j}\left(1-exp\left(-\frac{u_j x_j}{D}\right)\right)}\right)}\right) = \beta\left(\frac{Nl}{c+Nl}\right) \tag{S-15}$$

The critical length $L_c$ (Eq.S-13) provides an upper bound for the number of arms that can be used in the model

$$1 < \sqrt{\frac{c\beta}{D(N-1)}} \longrightarrow N < \frac{c\beta}{D} + 1 \tag{S-16}$$

To find the second polarization length $L_p$ we compare between the protrusive force in each arm when the global net flow is near zero (Eq.S-15), to the elastic force when the system is totally symmetric

$$\beta\left(\frac{L}{c+L}\right) = k\,(L-1) \tag{S-17}$$

where $L = \sum_j x_{tot}$.

By rearranging (Eq.S-17) we obtain

$$L_p = \frac{1}{2}(1-c) + \frac{\beta}{2k} + \sqrt{c + \left(\frac{1}{2}(1-c) + \frac{\beta}{2k}\right)^2} \tag{S-18}$$

For the moving case, we estimate the critical $\beta_c$ (denoted by $\beta_c^*$) by considering a smooth polarized motion along a linear track, with a steady-state treadmilling flow of $v = \beta - \frac{D}{c}$ [1]

$$\frac{dl}{dt} = \frac{1}{\Gamma}\left(v_{flow} - k(l-1)\right) \tag{S-19}$$

Under this estimation, we approximate that changes in the cell's arm lengths are small

$$\left|\frac{dl}{dt}\Gamma\right| \approx \eta < 1 \tag{S-20}$$

By combining (Eqs.S-19,S-20) we obtain the polarization length for the moving case

$$\eta = v_{flow} - k(L-1) \rightarrow L_p^* = \frac{c\,(c+\beta-\eta) - D}{ck} \tag{S-21}$$

and by equating $L_p^*$ (Eq.S-21) to $L_c$ (Eq.S-13) we obtain a closed form for $\beta_c^*$.

$$\beta_c^* = \frac{D}{2}\left(\frac{1}{c} + \frac{k}{\beta_c^* - \frac{D}{c} + k - \eta}\right) \tag{S-22}$$

### S-3. OSCILLATORY PATTERNS - EXTENDED ANALYSIS

In the simulations we find that two different mechanism govern the out-of-phase oscillations between the losing arms ($v_2$ and $v_3$). The first is discussed in the main text and in Fig.S-1 we present an extended analysis.

For the second mechanism we find that the solution branches of $v_2$ and $v_3$ are near the branching point after the symmetry breaking (Fig.S-2F-I). Since the length of $l_2$ and $l_3$ are nearly identical, they oscillate for a short period of time before $v_2$ and $v_3$ split (Fig.S-2C). During these small short oscillations in the ratios the solution branches of $v_2$ and $v_3$ flip (Fig.S-2F-I), and the system transitions between one branch and another until the difference between the losing arms is sufficiently large. We note that the same mechanism also occurs at the end of the time series when the cell escapes the junction, where $v_2$ and $v_3$ stabilize and converge into a single value. For this case, two solution branches converge into the branching point (end of the local flow time series in Fig.S-1C or S-8B).

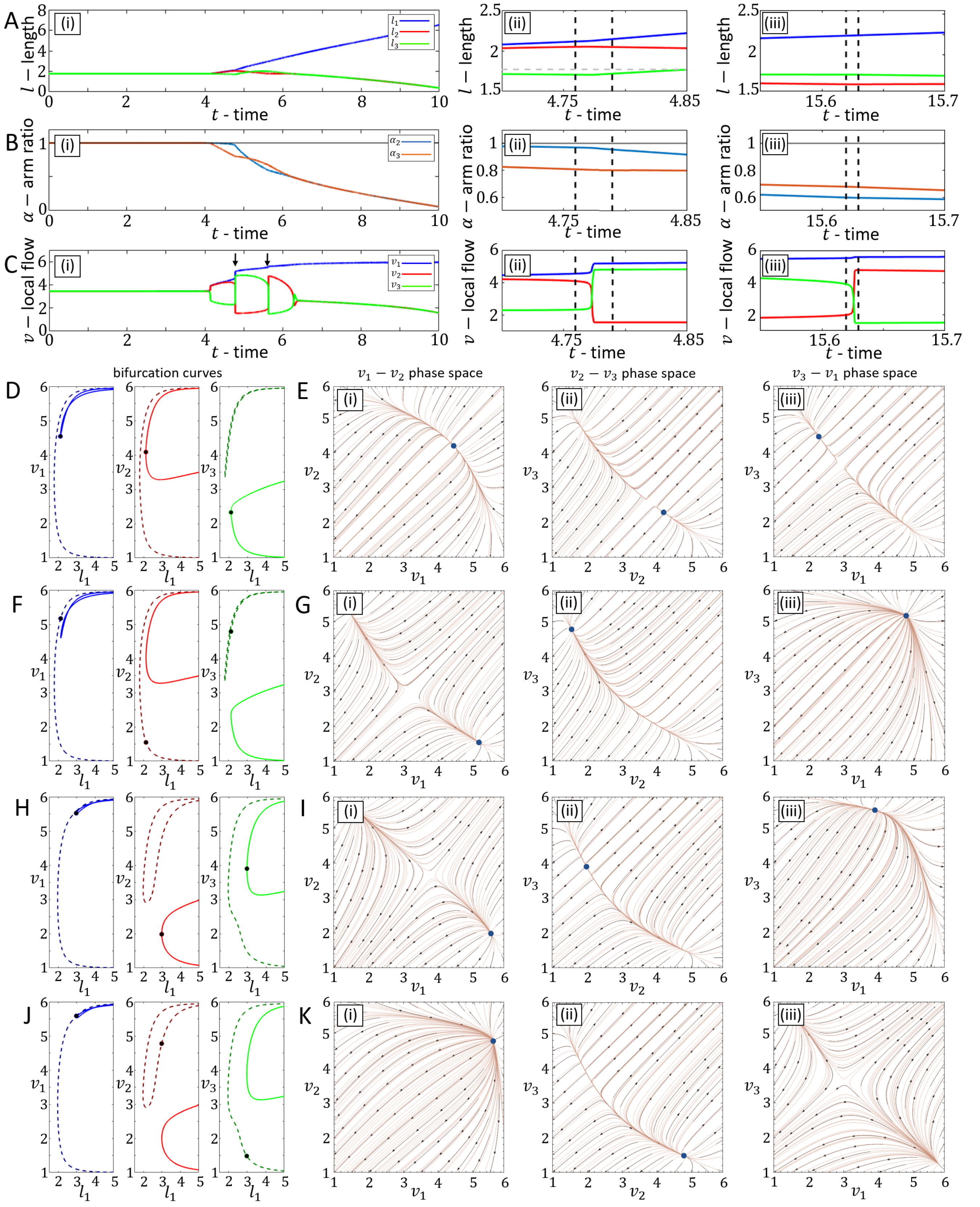


FIG. S-1: Analysis of the oscillation of type 1. A-C) Time series of the length, arm ratio and local flow respectively. (i) indicate the full time series. (ii) and (iii) zoom in the examined sections. Red/blue/Green curves in A and C indicate $v_1/v_2/v_3$. Blue/orange curves in B indicate $\alpha_1/\alpha_2$ respectively. Black dashed line indicates the examined cross-sections. D,F,G,J) The solution branches for $v_1$,$v_2$,$v_3$ (Blue,Red,Green) in the cross-sections. Points indicate the values from the simulation. E,G,I,K) Flow fields for (i) $v_2$-$v_1$, (ii) $v_3$-$v_2$ and (iii) $v_3$-$v_1$. Parameters: $\beta = 7.5$, $c = 3.85$, $D = 3.85$, $k = 0.8$, $f_s = 5$, $r = 8$, $\kappa = 20$, $\sigma = 10^{-7}$.

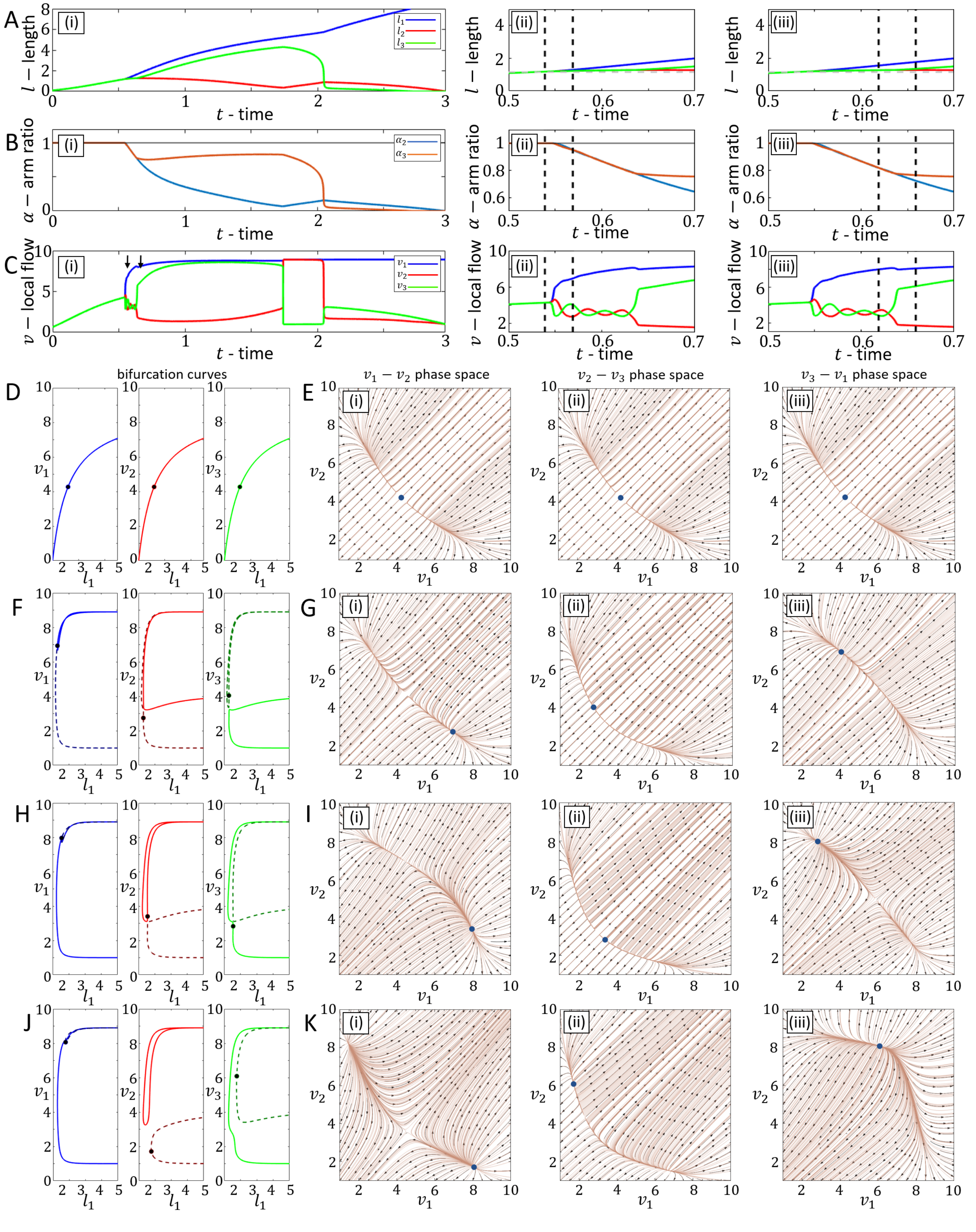


FIG. S-2: Analysis of the oscillation of type 2. A-C) Time series of the length, arm ratio and local flow respectively. (i) indicate the full time series. (ii) and (iii) zoom in the examined sections. Red/blue/Green curves in A and C indicate $v_1/v_2/v_3$. Blue/orange curves in B indicate $\alpha_1/\alpha_2$ respectively. Black dashed line indicates the examined cross-sections. D,F,G,J) The solution branches for $v_1$,$v_2$,$v_3$ (Blue,Red,Green) in the cross-sections. Points indicate the values from the simulation. E,G,I,K) Flow fields for (i) $v_2$-$v_1$, (ii) $v_3$-$v_2$ and (iii) $v_3$-$v_1$. Parameters: $\beta = 8.5$, $c = 3.85$, $D = 3.85$, $k = 0.8$, $f_s = 5$, $r = 8$, $\kappa = 20$, $\sigma = 10^{-7}$.

### S-4. EXPERIMENTAL METHODS

#### Glioma cells

**NNI-21 cell line:** hGPC NNI-21, were originally isolated in the laboratory of Carol Tang at the National Neuroscience Institute (Singapore) from GBM tumor specimens obtained with informed consent and de-identified in accordance with the SingHealth Centralised Institutional review Board A. This cell line was previously used and published [2]. The hGPCs were maintained, as tumorspheres, in DMEM/F12, supplemented with sodium pyruvate, non –essential amino acid, glutamine, penicillin/streptomycin, B27 supplement, bFGF (20 [ng/ml]), EGF (20 [ng/ml]), and heparin (5 [µg/ml]).

**Transfection:** For transfection and migration assays, NNI-21 cells were cultured as monolayers on laminin (10 [µg/ml]) coated petri dishes for 3-5 [days] before transfection. NNI-21 transfections were performed with a Neon electroporator (Invitrogen) as per manufacturer's recommendations as follow: 10 [µg] DNA for 1,000,000 cells, in 100 [µl] buffer R were electroporated at 1600 [V], 20 [ms], 1 choc. GFP-actin was from E. Lemichez (Pasteur Institute, France).

**Micropatterning:** Honey comb-shaped micropatterns (7 [µm] width, 150 [µm] gap) were obtained using deep UV lithography [3, 4]. Chrome masks were designed and produced by Photomask portal (Richardson, TX 75082). Briefly, coverslips were plasma cleaned for 3 [min], incubated in 0.1 [mg/ml] PLL-g-PEG in 10 [mM] HEPES, 7.4 [ph], at room temperature for 1 [hr], and put in contact with the chrome mask using a water drop (3 [µl]). The sandwich was then placed under UV 180 [nm] for 10 [min] to oxidize the PLL-g-PEG under the transparent areas. The coverslips were then mounted in imaging chambers and incubated with a solution of laminin (10 [µg/ml] in PBS) for at least 1 [hr] at 37°C, rinsed in PBS and medium before cell seeding.

**Imaging:** Long term imaging, in Phase contrast microscopy was performed on a leicaAM TIRF MC system with a 10X objective (Leica HCX PL FLUOTAR 10x/0.30NA PH1 Objective). The microscope was controlled by Leica Application Suite AF software and images were acquired with an Andor iXon DU-8285-VP camera (1 image/30 [sec]). GFP-actin was imaged with a Thunder Imaging System (Leica) based on a Leica DMi8 microscope equipped with a Leica DFC9000 GT sCMOS camera. The images were acquired with a 40X objective (HC PL APO 40X 0.95NA) using Leica LAS X software (1 image/2 [min]). Both microscopes were equipped with temperature, humidity, and CO2 control.

#### HUVEC cells

**HUVEC cell line:** Corning culture dishes with 100 [mm] diameter were coated with fibronectin from bovine plasma (1 [µL/mL]) for 20 [min] at 37 °C and 5 % CO2. Afterwards, dished were washed once with DPBS (1X, Gibco). The HUVECs were cultured in ready-to-use full endothelial growth medium that included the Basal Medium and the SupplementMix (PromoCell) and was supplemented with 1 % Penicillin-Streptomycin (PenStrep, ThermoFisher). HUVECs were incubated at 37 °C and 5 % CO2 for 2 [days] until a monolayer was reached to then passage them again. Every passage was done by detaching with 3 [mL] of TrypLE Express (with Phenol Red, 1X, Gibco, ThermoFisher) for 2 [min] and then plating 1,5x106/[mL] in a new coated Corning culture dishes with 100 [mm] diameter in a total of 10 [mL] new full endothelial growth medium. Primary Human Umbilical Vein Endothelial Cells isolated from the vein of the umbilical cord of single, pooled donors (HUVECs) were used between passages 5 and 12.

**Transfection:** HUVECs were cultured until reaching a confluence of 80% and then were detached as mentioned for the passage. Then transfected, using the 4D-Nucleofector™ X Unit (Lonza ),with 1 [µg] of F-tractin GFP plasmid and the P5 Primary Cell 4D-Nucleofector™ X Kit to then electroporate corresponding to the transfection program(CA-167) to then keep overnight (around 16 [hr]) in culture previous to use.

**Micropatterning:** For HUVECs Photopatterning was performed using a Digital Micromirror Device (Primo™, ALVEOLE) coupled to a T2i Eclipse microscope (Nikon). A Polydimethylsiloxan (PDMS) stencil with a circle area of 5 [mm] was placed inside a 35 [mm] diameter glass bottom FluoroDish (World Precision Instruments) and cleaned with a PlasmaCleaner (PDC-32G-2, Harrick Plasma) for 5 [min]. Afterwards, the area within the stencil was coated with 0.1 [mg/mL] pLL-PEG (SuSoS, Surface Technology) for 1 [hr] at room temperature. The remaining pLL-PEG was washed out with sterile water. After that the photo-activator (PLPP™, ALVEOLE) was added and then degraded under UV illumination at a power of 600 [mJ/mm2]. Hexagonal patterns (width 20 [µm], side length of 140 [µm]) were created using the software Inkscape 1.0.2-2. The samples were washed with water to remove the PLPP and coated for 20 min with fibronectin (1 [µL/mL]) mixed with fibrinogen (1:10) to visualize the patterns and then rinsed with 1X DPBS. Then, 10,000 cells were plated and incubated at 37°C and 5 % CO2 overnight (around 16 [hr]) to allow their adhesion. Once attached samples were carefully washed with medium.

**Imaging:** Live cell imaging was performed with a Leica Dmi8 inverted microscope equipped with a APO 10x/0.45 PH1, and APO 40x/0.95 objectives. Images were recorded with an ORCA-Flash4.0 Digital camera (Hamamatsu Photonics) using the MetaMorph Version 7.10.3.279 software (Molecular Device). The acquisition for movies at low magnification or actin visualization was 3 [min], and for high magnification 30 [sec]. During every acquisition, cells were kept under optimal conditions at 37°C and 5 % CO2 because of a chamber lid that was placed on the microscope stage.

### S-5. EXTENDED COMPARISONS BETWEEN THEORY AND EXPERIMENTS

In this section we display further examples of hGPC and HUVEC experiments, and their comparison with simulations. In Figs.S-3,S-4 we provide more examples of HUVEC and hGPC cells which perform length oscillations, as well as out-of-phase oscillations in the actin activity at the tips. In Fig.S-4A-D the hGPC cell performs a stick slip event which locates the cell on the center of the junction when it is short and un-polarized, which allows us to observe the symmetric growth of all the protruding arms prior to the symmetry breaking.

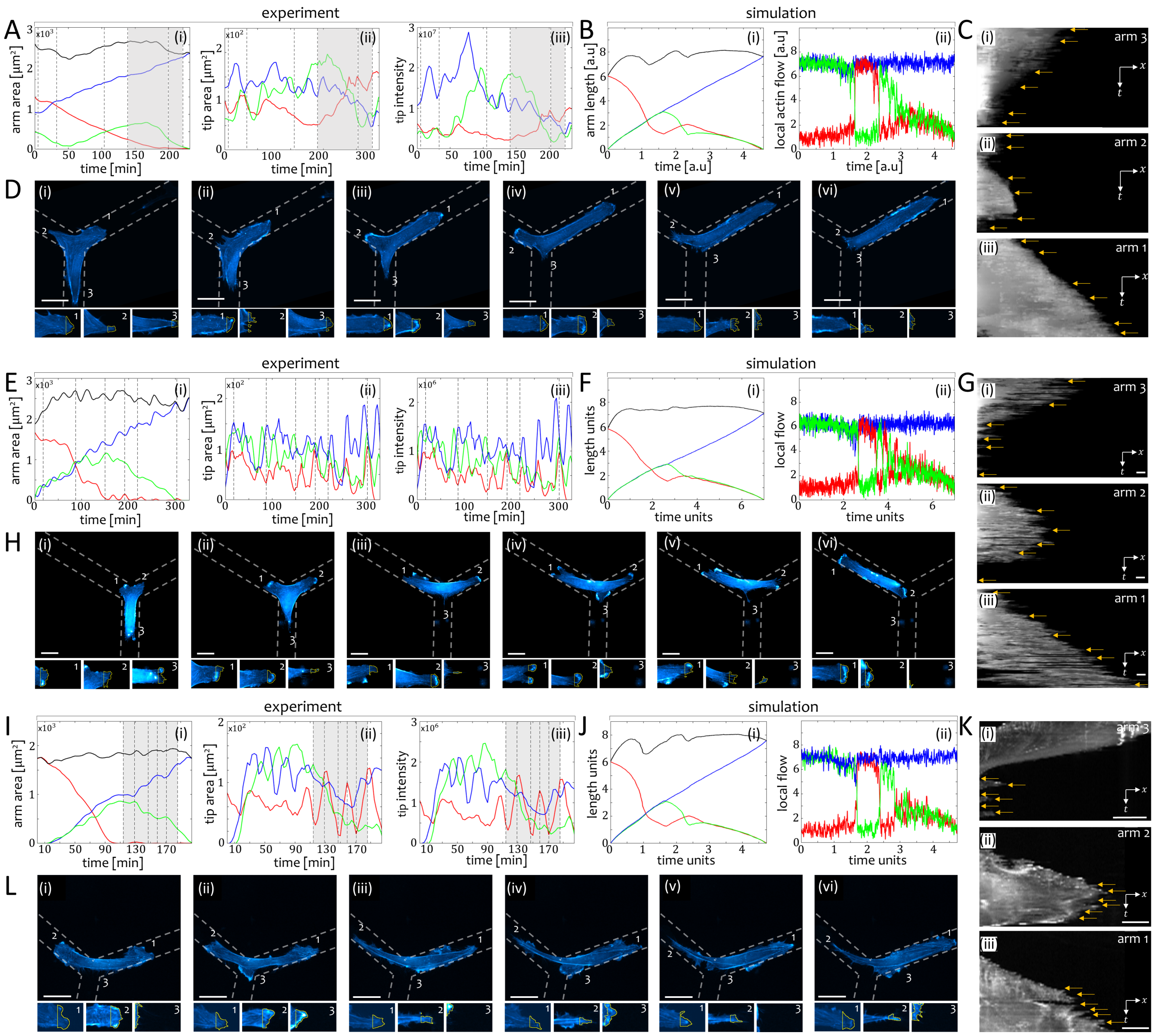


FIG. S-3: Comparison between HUVEC cell experiments and simulations. A) i) Total area of the arms. ii) Lamellipodia (arm tip) area. iii) Lamellipodia actin intensity. B) Simulation with $(\beta, \sigma) = (6.8, 0.5)$. i) Arm length. ii) local actin flows. Blue/Green/Red indicate arm 1/2/3. Black indicate the total cell length. Gray section indicate the regions where the actin flow flips between the losing arms. Gray dashed lines correspond to the images in D and the arrow markings in C. C) Arm kymographs. D) Time-lapse images (movie S1). Upper panels: Whole cell. Lower panels: Enlarged arm protrusions. Yellow is a 10 [μm] window from which the intensity was measured. E-H) Same as A-D with $(\beta, \sigma) = (6.5, 0.6)$. Time-lapse images correspond to movie S2. I-L) Same as A-D with $(\beta, \sigma) = (7, 0.6)$ used in the simulation. Time-lapse images correspond to movie S3. Scale bars in (D,H,L) are 100 [μm]. Other simulation parameters: $c = 3.85$, $D = 3.85$, $k = 0.8$, $f_s = 5$, $r = 5$, $\kappa = 20$, $\delta = 250$.

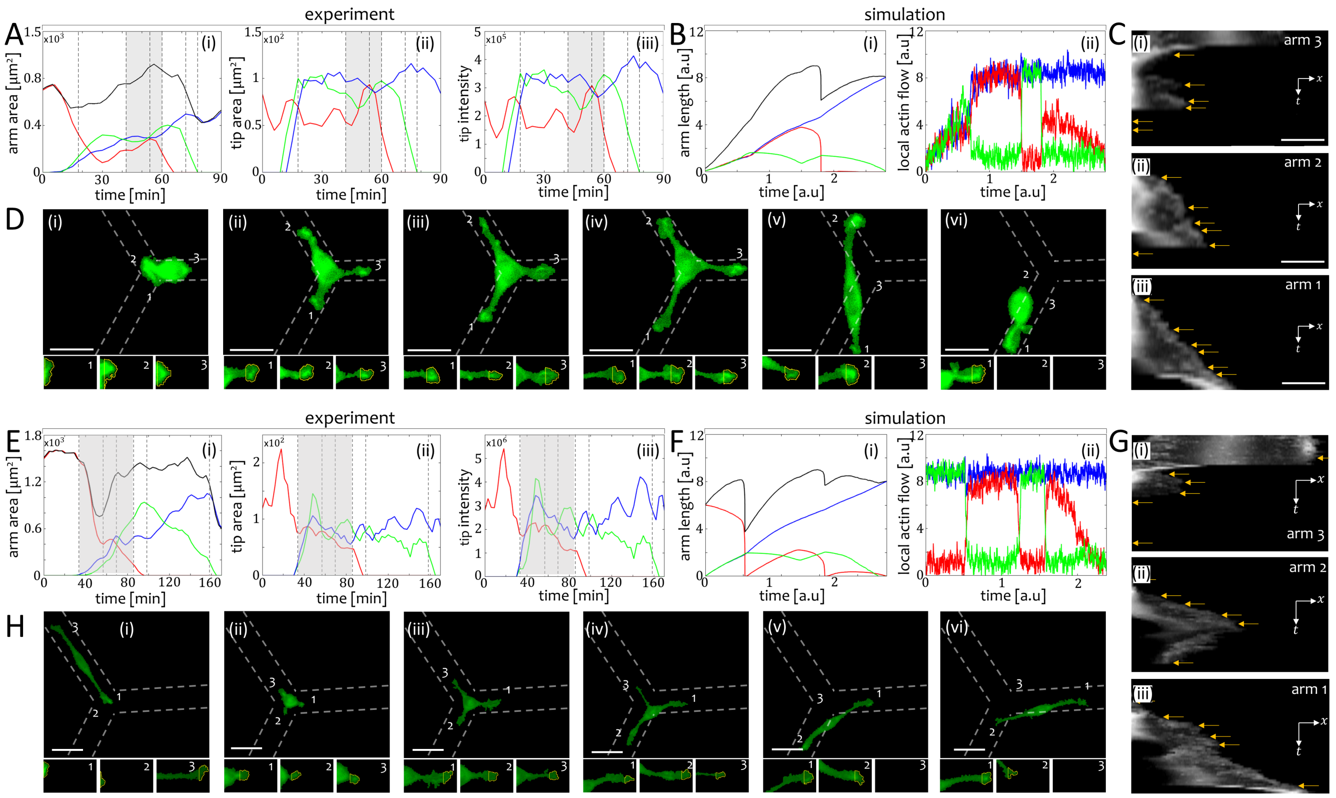


FIG. S-4: Comparison between hGPC cell experiments and simulations. A) i) Total area of the arms. ii) Lamellipodia (arm tip) area. iii) Lamellipodia actin intensity. B) Simulation with $(\beta, \sigma) = (8.5, 0.7)$. i) Arm length. ii) local actin flows. Blue/Green/Red indicate arm 1/2/3. Black indicate the total cell length. Gray section indicate the regions where the actin flow flips between the losing arms. Gray dashed lines correspond to the images in D and the arrow markings in C. C) Arm kymographs. D) Time-lapse images (movie S4). Upper panels: Whole cell. Lower panels: Enlarged arm protrusions. Yellow is a 10 [µm] window from which the intensity was measured. E-H) Same as A-D with $(\beta, \sigma) = (8.7, 0.7)$. Time-lapse images correspond to movie S5. Scale bars in (D,H) are 100 [µm]. Other simulation parameters: $c = 3.85$, $D = 3.85$, $k = 0.8$, $f_s = 5$, $r = 5$, $\kappa = 20$, $\delta = 250$.

In Fig.S-5 we show more examples of HUVEC cell experiments which perform length oscillations. Fig.S-5E-G shows a cell which enters the junctions, and turns back in the arm from which it entered the junction, a results which is found when the noise level is sufficiently large (Fig.6A in the main text).

In Fig.S-6 we provide two example sets of HUVEC cells which perform similar dynamics: One set has few stick-slip events (Fig.71S-6A), and the other set has more stick-slip events, while the corresponding escape times are roughly doubled (Fig.S-6C). These examples correspond to the difference in simulations with low and high noise amplitudes, irrespective to the value of $\beta$ (Fig.S-6B,D, and Fig.6 in the main text).

### S-6. ESCAPE TIME - EXTENDED ANALYSIS

This section extends the escape time analysis to cases where the actin flow instantaneously returns to its steady-state value, $\delta \to 0$ (Fig.S-7). The analysis is performed for the symmetric case.

Here, we divide the dynamical behavior as a function of $\beta$ into three regions, with respect to the mean escape time and length. In region 1 the mean escape time/length decreases/increases smoothly as $\beta$ increases (Fig.S-7A,B,E). In this region we find that prior to the escape, the two losing arms compete, and exhibit oscillations in the local flows ($v_i$) while the motion of the arms is smooth (Fig.S-8A,B), regardless of the value of $\delta$. Region 2 is characterized by a decrease in the mean escape length as $\beta$ increases (Fig.S-7B), which can be attributed to a stick-slip event which occurs in one of the losing arms during the escape (Fig.S-8C,D). In this region the escape time and escape length variances demonstrate an irregular behavior (Fig.S-7C,D), mostly for $\delta \to \infty$. This irregularity is attributed to a series of short oscillations between the losing arms which occur directly after the symmetry breaking (panels (*) below

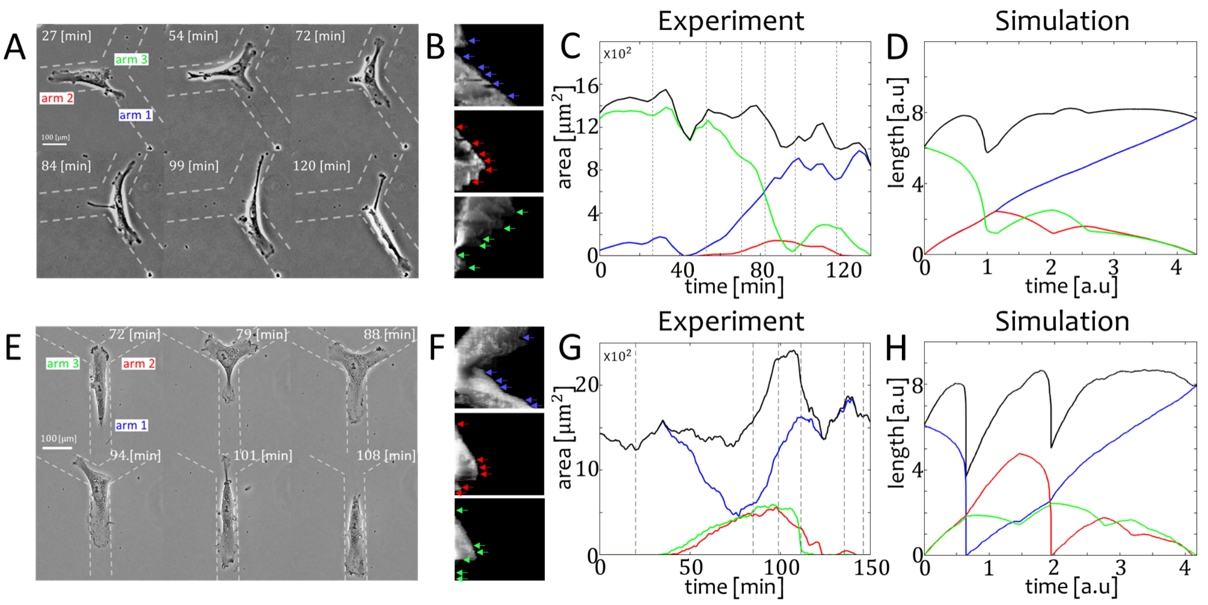


FIG. S-5: Comparison between HUVEC experiments and simulations. A) Time-lapse images (movie S6). B) Arm kymograph. Blue/Red/Green arrows indicate the time stamps which correspond to the images in A. C) Time series of the areas of the arms. vertical dashed lines correspond to the images in A. D) Simulation time series of the arm lengths with $(\beta, \sigma) = (7.2, 0.3)$. Blue/Red/Green indicate arm 1/2/3. Black indicates the total length of the cell. E-H) Same as A-D. Simulation correpond to $(\beta, \sigma) = (8.0, 1.2)$. Time-lapse images correspond to movie S7. Other simulation parameters: $c = 3.85, D = 3.85, k = 0.8, f_s = 5, r = 5, \kappa = 20, \delta = 250$.

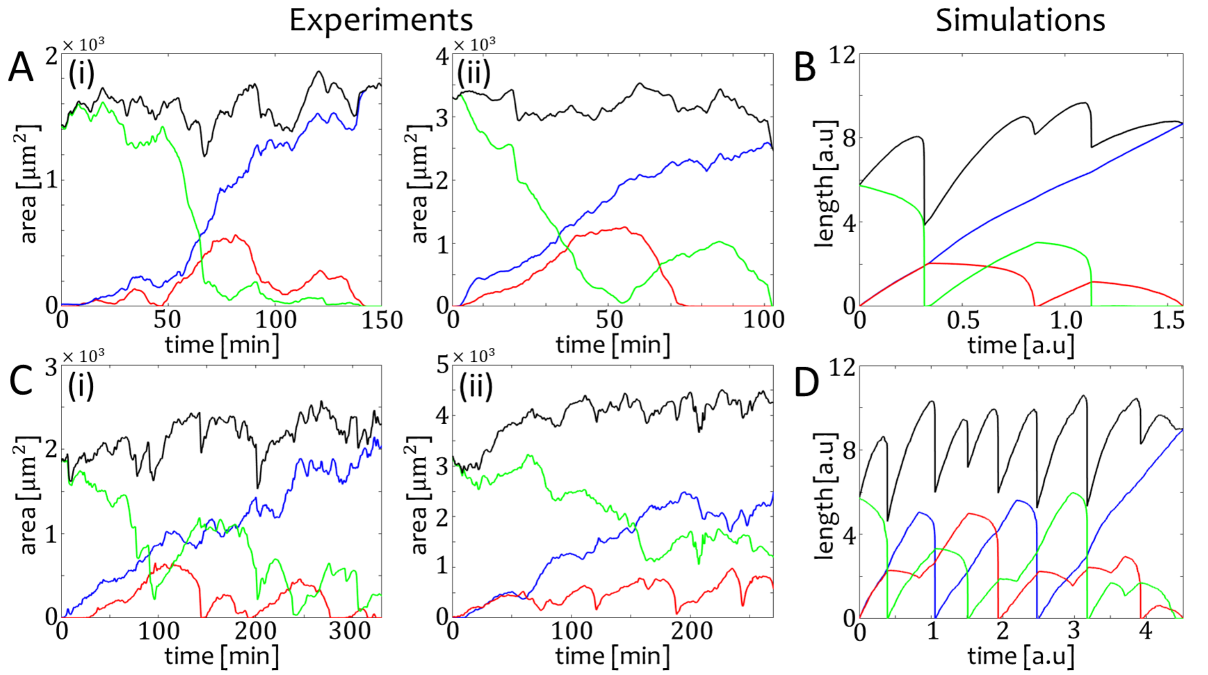


FIG. S-6: Comparison between HUVEC cell experiments and simulations with different noise amplitudes. A) Experiment. Time series of the arm area for two cells (i-ii) which demonstrate single or few stick-slip events. B) Simulation with $\beta = 11$ and a low amplitude of noise $\sigma = 0.5$. C) Experiment. Time series of the arm area for two cells (i-ii) which demonstrate single or few stick-slip events. D) Simulation with $\beta = 11$ and a high amplitude of noise $\sigma = 2.3$. Blue/Red/Green indicate arm 1/2/3. Black indicates the total length. Other simulation parameters: $c = 3.85, D = 3.85, k = 0.8, f_s = 5, r = 5, \kappa = 20, \delta = 250$.

Fig.S-7C). In this region, the system is highly sensitive, and the escape time depends on the number of oscillations. In region 3 the dynamics for $\delta \to \infty$ are characterized by a symmetry breaking with no oscillations in the flows (Fig.S-8E), while for $\delta \to \delta_0$ we find a bi-stability between a slow and a fast process (Fig.S-7A,B,G and Fig.S-8F,G).

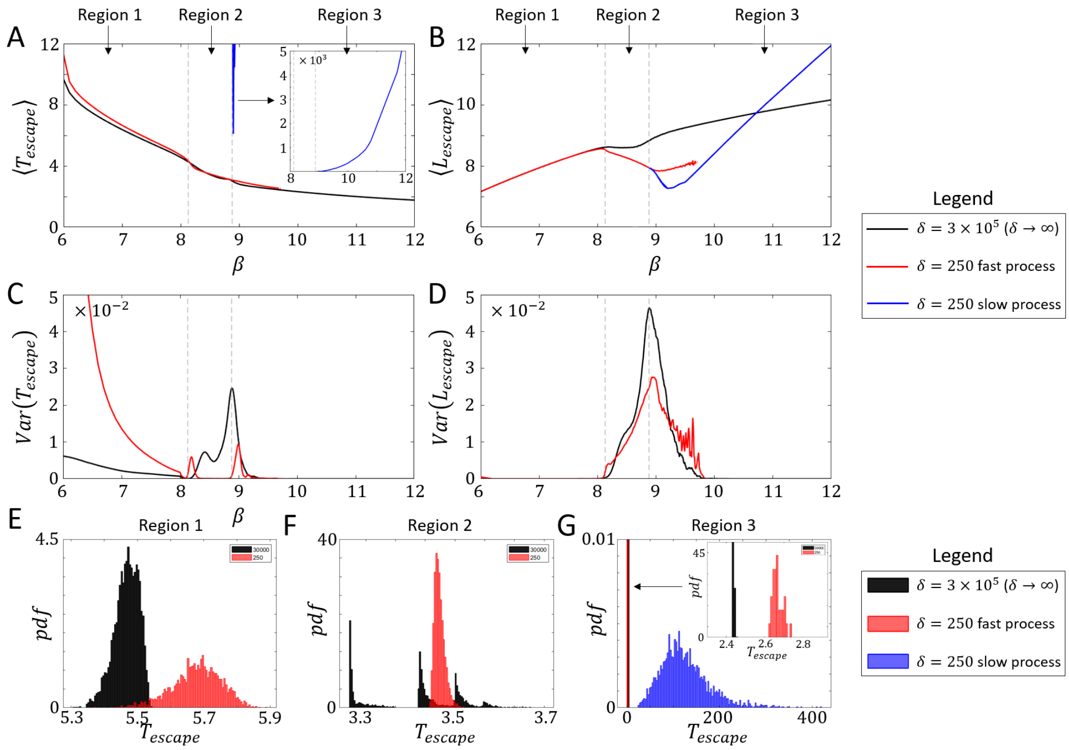


FIG. S-7: A) The mean escape time as a function of $\beta$. Right upper inset indicates the slow process. B) The escape length as a function of $\beta$. C) The variance of the escape time as a function of $\beta$. D) The variance of the escape length as a function of $\beta$. E-F) Examples of typical escape time histograms from region 1 ($\beta = 7.5$), region 2 ($\beta = 8.5$), and region 3 ($\beta = 9.5$). Black stands for $\delta = 30000$, and Red/Blue stands for the fast/slow process of $\delta = 250$. Parameters: $c = 3.85$, $D = 3.85$, $k = 0.8$, $f_s = 5$, $r = 8$, $\kappa = 20$, $\sigma = 10^{-7}$.

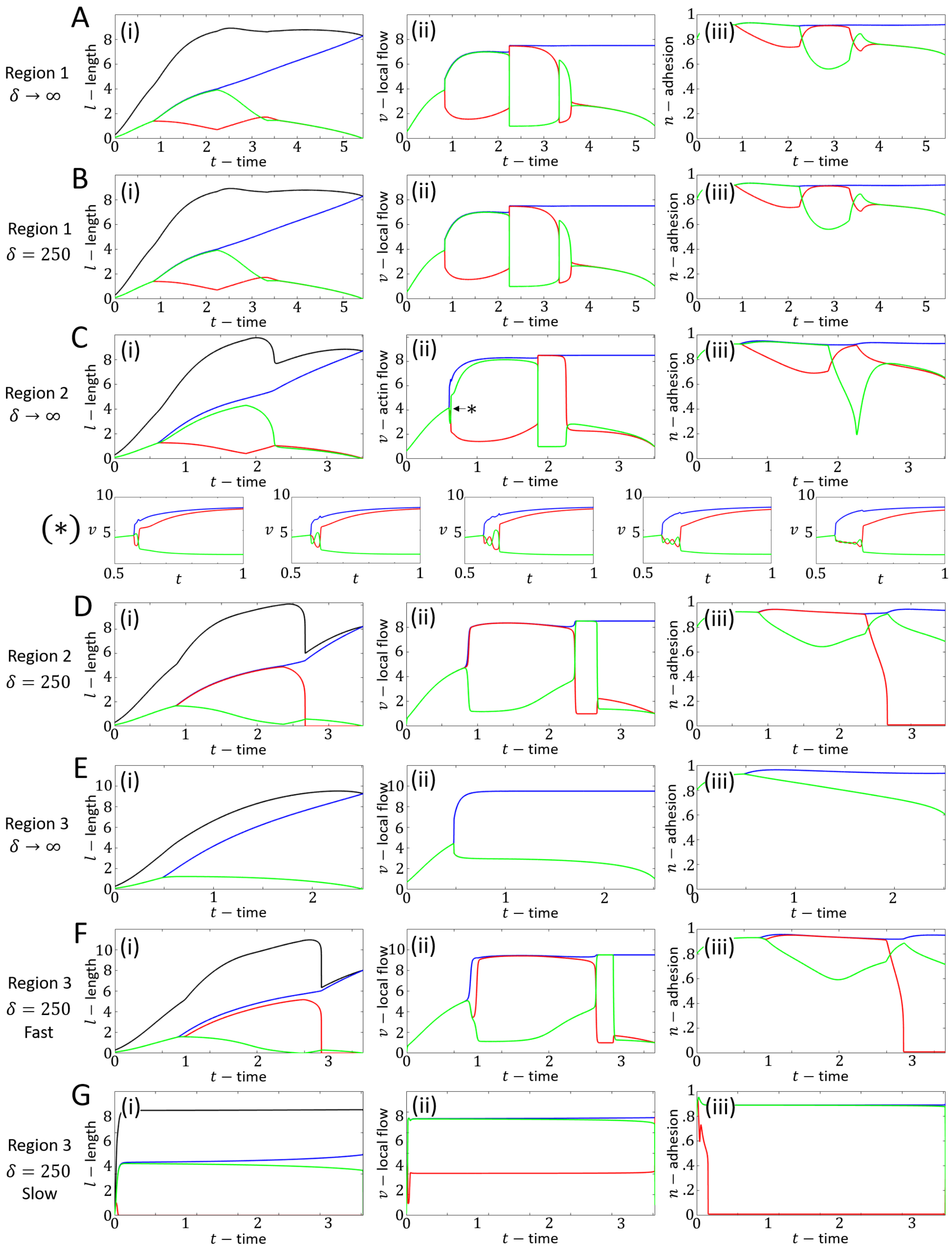


FIG. S-8: A-D) Examples of trajectories from the simulation for $\delta = 30000$ and $\delta = 250$: A-B) Region 1. C-D) Region 2. E-G) Region 3. Panel (i): The length. Blue/Red/Green stand for arm 1/2/3 respectively. Black stands for the total length. Panel (ii): The local flow. Blue/Red/Green stand for arm 1/2/3 respectively. Panel (iii): The adhesion concentration. Blue/Red/Green stand for arm 1/2/3 respectively.

### S-7. THE SLOW PROCESS WITH DIFFERENT FINITE $\delta$ VALUES

In this section we further analyze the slow process dynamics. We examine intermediate values of $\delta$ ($250 < \delta < 3\times10^5$), where for the analysis we choose $\delta = 9000$). We find that there are several escape times for the slow process (Fig.S-9A,B). The fast process also has several different escape times however with a very small time margin between them (Fig.S-9C).

In the $v_1$-$v_2$ phase-space we find that the reason behind the splitting in the escape time lies with the oscillations of the second type (Fig.S-8) which occur right after the symmetry breaking. The number of oscillations in the local flows of the losing arms will determine the escape time. As the number of these oscillations increase the escape time increases as well. The $v_1$-$v_2$ phase-space also allows us to better understand the mechanism behind the slow process, where for a reason which is not yet understood the trajectory of $v_1$-$v_2$ in the phase-space for the slow process climbs back slowly to the local minimum (Fig.S-9E,F), before the local flows complete a full oscillation (Fig.S-9D)

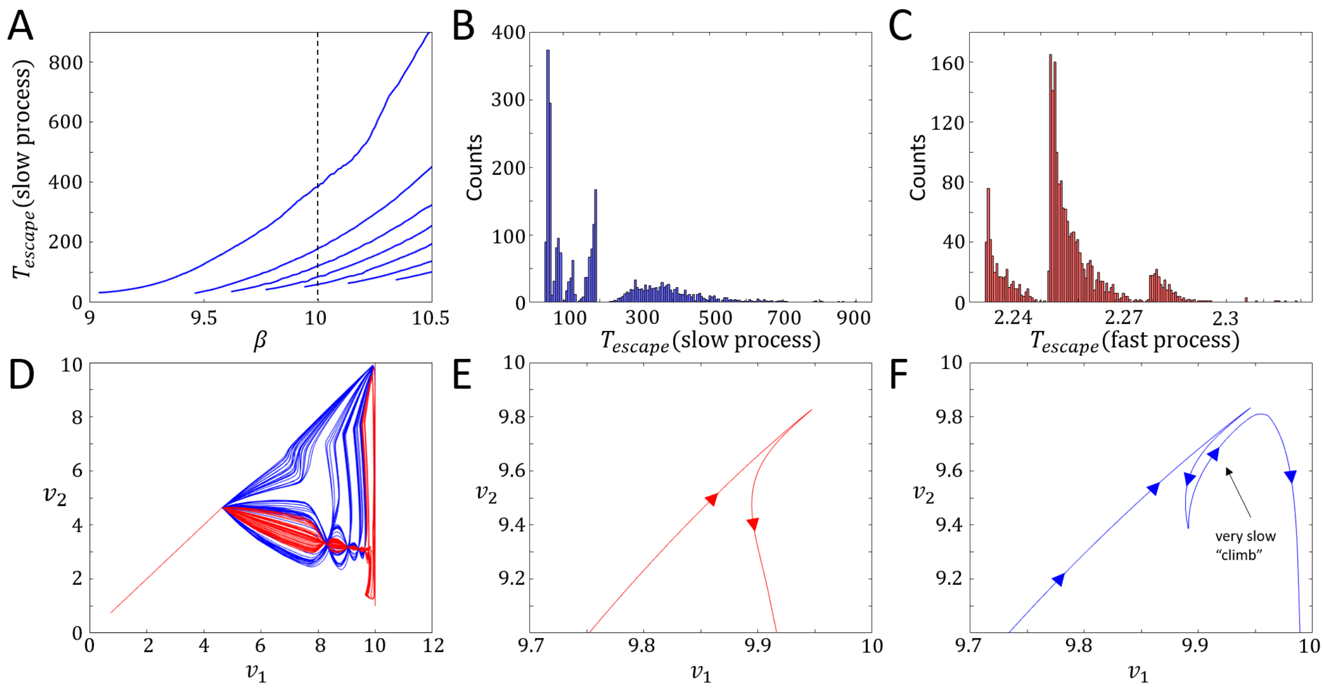


FIG. S-9: A) The mean escape time for the slow processes with $\delta = 9000$. Black dashed line indicate the analyzed cross-section of $\beta = 10$. B-C) Histograms of the escape times for the slow/fast processes (blue/red) respectively. D) $v_1$-$v_2$ phase-space trajectories of 200 simulations. Red/Blue curves indicate the fast/slow processes. E-F) Zoom in on the $v_1$-$v_2$ phase-space dynamics of both the fast (E) and the slow (F) processes in the proximity of the local saddle point, indicating the difference in the dynamics. Parameters: $\beta = 10$, $c = 3.85$, $D = 3.85$, $k = 0.8$, $f_s = 5$, $r = 8$, $\kappa = 20$, $\delta = 9000$, $\sigma = 10^{-7}$.

### S-8. NOISE EFFECTS ON THE ESCAPE TIME AND DYNAMICS AT THE JUNCTION

In this section we provide the full noise influences analysis of the moving case. Here, we examine three value of $\beta$, which characterize regimes of smooth and stick-slip motion for the cell along a linear track [1], and compare between the effects of large and small amplitudes of noise on the trajectories. In the low $\beta$ regime ($\beta = 6$), where the motion of the arms are smooth (Fig.S-10A,D,G,H), we find that the cell will escape the junction with probability $p = 1$ for low noise (Fig.S-10A,D(i),G). As the noise amplitude increases the probability to escape decreases. For large noise amplitudes the cell remains trapped at the junction with zero probability to escape, as the actin flow can not stabilize its polarity due to the large fluctuations (Fig.S-10A,D(ii),H). In the intermediate $\beta$ regime ($\beta = 10$), we find that the cell escapes the junction with probability $p = 1$ for all noise amplitudes examined for this section (Fig.S-10B). For low values of noise the cell performs a single stick-slip event in one of the losing arms (Fig.S-10I), while for larger amplitude of noise the cell can perform more stick-slip events (Fig.S-10J), and the parity of the stick-slip events in the losing arms determines if the cell will reverse its motion and leave the junction along the arm from which it has arrived (Fig.S-10E(ii)). We note that as the noise level increase (beyond the values presented in Fig.S-10B), the protruding arms of the cell can perform more stick-slip cycles before escaping the junction, and when the amplitude of the noise is sufficiently large, the cell will remains trapped on the junction, as exhibited in the low $\beta$ regime (Fig.S-10A). In the regime of large $\beta$ ($\beta = 14$), we find that large values of noise push the system away from the meta-stable state of the slow process which is observed in the deterministic system (Fig.S-10C) and prevent the cell from being trapped on the junction, while for low values of noise the system maintains its bi-stable behavior between a slow and a fast process (Fig.S-10F,K,L). We also observe that the total cell length elongates significantly while the cell is trapped in the junction (Fig.S-10H,L), compared to the length of the cell when it is moving along a linear track [1]. This means that after the cell escapes the junction, the length of the cell shrinks rapidly before it moves away along a linear path.

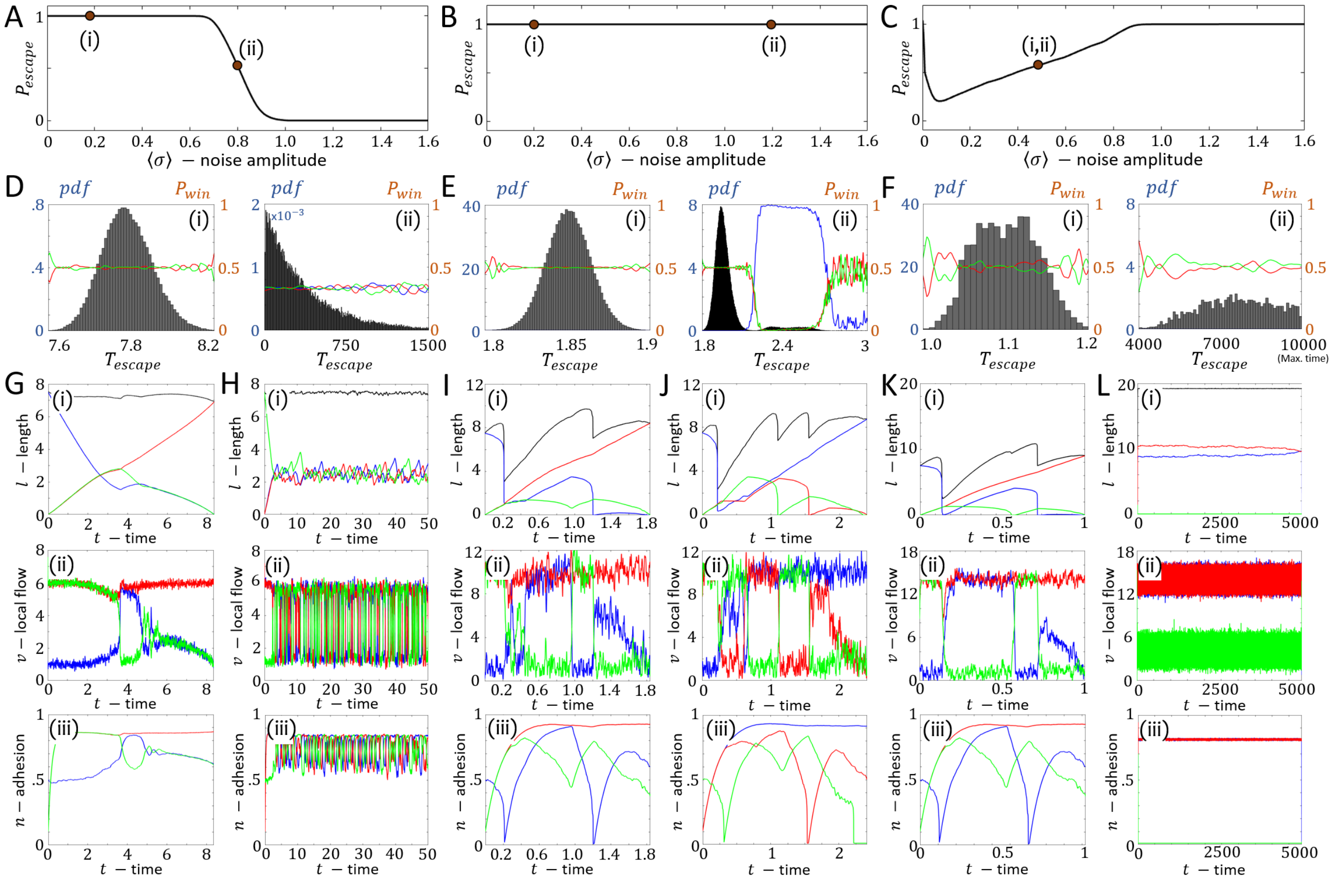

FIG. S-10: A-C) The probability to escape as a function of the noise amplitude $\sigma$ for $\beta = 6, 10, 14$ respectively. D-F) The escape time probability density function (left y-axis in blue) and the probability for each arm to win (right y-axis in orange) for $\beta = 6, 10, 14$. Blue/Red/Green indicate arm 1/2/3. (i-ii) relate to $\sigma$ values of the orange points in panels A-C. G-L) Examples of times series of the length (i), the local actin flow (ii), and the adhesion concentration (iii), for $\beta = 6$, $\sigma = 0.2$ (G), $\beta = 6$, $\sigma = 0.8$ (H), $\beta = 10$, $\sigma = 0.8$ for a single stick-slip event (I), $\beta = 10$, $\sigma = 0.8$ for two stick-slip events (J), $\beta = 14$, $\sigma = 0.5$ for a fast process, and $\beta = 14$, $\sigma = 0.5$ for a slow process. Parameters: $c = 3.85, D = 3.85, k = 0.8, f_s = 5, r = 5, \kappa = 20, \delta = 250$.

### S-9. SUPPLEMENTARY MOVIES

**Movie 1:** Human glioma propagating cells (hGPCs) migrating in a mouse brain slice (Fig.1A).

**Movie 2:** Example of an hGPC migrating along a laminin-coated Y-junction (Fig.1B).

**Movie 3:** Example of a HUVEC migrating along a fibronectin-coated Y-junction (Fig.1C).

**Movie 4:** HUVEC experiment showing out-of-phase oscillations in arm length and actin activity (Fig.4D).

**Movie 5:** hGPC experiment showing out-of-phase oscillations in arm length and actin activity (Fig.4H).

**Movie 6:** hGPC migrating over a blood vessel bifurcation (Fig.4I).

**Movie 7:** hGPC experiment, where two competing arms extend, as in the ”slow process” (Fig.5H).

**Movie 8:** hGPC experiment, where the cell is not-deciding on a new direction of motion (Fig.6C).

**Movie 9:** HUVEC experiment, where the cell is not-deciding on a new direction of motion (Fig.6F).

**Movie S1:** HUVEC experiment showing out-of-phase oscillations in arm length and actin activity (Fig.S3D).

**Movie S2:** HUVEC experiment showing out-of-phase oscillations in arm length and actin activity (Fig.S3H).

**Movie S3:** HUVEC experiment showing out-of-phase oscillations in arm length and actin activity (Fig.S3L).

**Movie S4:** hGPC experiment showing an un-polarized cell which resides in the center of the junction (after a stick-slip event), and performs a symmetric spreading in all the protruding arms (Fig.S4D).

**Movie S5:** hGPC experiment showing out-of-phase oscillations in arm length and actin activity (Fig.S4H).

**Movie S6:** HUVEC experiment showing a normal escape pattern (Fig.S5A).

**Movie S7:** HUVEC experiment showing a cell which ”reflects” back in the direction from which it has entered the junction (Fig.S5E).

All scale bars are 100 [µm].

---

[1] J. E. Ron, P. Monzo, N. C. Gauthier, R. Voituriez, and N. S. Gov, Physical Review Research **2**, 033237 (2020).
[2] P. Monzo, M. Crestani, Y. K. Chong, A. Ghisleni, K. Hennig, Q. Li, N. Kakogiannos, M. Giannotta, C. Richichi, T. Dini, et al., Developmental cell (2021).
[3] A. Azioune, N. Carpi, Q. Tseng, M. Thery, and M. Piel, in *Methods in cell biology* (Elsevier, 2010), vol. 97, pp. 133–146.
[4] M. Crestani, T. Dini, N. C. Gauthier, and P. Monzo, STAR protocols **3**, 101331 (2022).
